# Therapeutic antisense oligonucleotide mitigates retinal dysfunction in a pig model of CLN3 Batten disease

**DOI:** 10.1101/2025.05.30.656864

**Authors:** Matthew P. Stratton, Jessica L. Centa, Vicki J. Swier, Wanda L. Pfeifer, Clarissa D. Booth, Karlee Albert, John L. Hunyara, Mitchell J. Rechtzigel, Fox J. Duelli, Hannah G. Leppert, Frank Rigo, Trisha Smit, Paymaan Jafar-Nejad, Jill M. Weimer, Arlene V. Drack, Michelle L. Hastings

## Abstract

CLN3 Batten disease is a lethal pediatric autosomal recessive neurodegenerative disease caused by mutations in the *CLN3* gene. Typically, the disease manifests as vision loss early in life and progresses to neurological dysfunction and death in young adulthood. Therapeutic development has focused on treating the central nervous system. However, such therapies may not protect against vision loss, which has a significant impact on quality of life. We have shown that a splice-switching antisense oligonucleotide (ASO) delivered to the central nervous system can reduce neurological disease burden in mouse models of CLN3 disease. Here, we report on a similar ASO approach for treating CLN3 Batten disease retinal dysfunction in a pig model of the disease, which is more representative of human vision. A single intravitreal injection of ASO induces robust exon skipping in the retina for up to 12 months. The ASO treatment resulted in higher amplitudes on electroretinograms, suggesting mitigation of retinal dysfunction at early timepoints of disease. One ASO that efficiently induces exon skipping *in vivo* was well-tolerated and targets a region of *CLN3* that is conserved in humans, making it a promising candidate for treating the disease in humans. Our findings demonstrate the potential utility of an ASO-based approach to treat retinal dysfunction in CLN3 Batten disease and generally supports the use of ASOs for treating eye diseases.

**One Sentence Summary:** Splice-switching antisense oligonucleotides delivered by intravitreal injection are safe and show efficacy in preventing early retinal dysfunction in a pig model of CLN3 Batten disease.

## INTRODUCTION

CLN3 Batten disease, also known as CLN3 disease or juvenile neuronal ceroid lipofuscinosis (JNCL), is an autosomal recessive lysosomal storage disorder, affecting as many as 1 in 25,000 people in some populations (*1–3*). In classical presentations of the disease, children develop normally until symptom onset at 4-8 years of age when vision loss begins, typically as the first clinical sign of the disease (*4*). Disease symptoms advance during adolescence to include seizures, difficulty speaking, loss of motor function, cognitive decline, psychiatric problems, and ultimately premature death in the second or third decade of life (*5–8*). There are currently no effective treatments for CLN3 disease.

Vision loss causes a significant reduction in the quality of life for people with CLN3 disease, making it a therapeutic priority at early stages of the disease (*4*). In children with CLN3 disease, visual acuity is mildly affected initially, but vision loss due to retinal degeneration worsens rapidly, progressing from normal vision to functional blindness over 24-36 months, although light-dark perception may be retained for several years (*6, 9*). Electroretinograms (ERG) of patients with early CLN3 disease are electronegative (*9–11*), but as the disease progresses, the ERG is severely reduced (*12*) with no rod response (*9, 11, 13*), diminished and delayed cone responses (*9, 13*), reduction in the a-wave response (*10, 11*), even greater loss of b-wave amplitudes (*11*), reduction in the b:a ratio (*9*), and are ultimately completely undetectable (*9, 10*).

*CLN3* is a 15-exon gene encoding a transmembrane protein of unknown function that localizes to the membrane of lysosomes and endosomes via two lysosomal targeting sequences (LTS) (*14–17*). The most common disease-causing variant in *CLN3*, accounting for ∼85% of disease alleles, is a 966 base pair deletion encompassing the entirety of exons 7 and 8 (*CLN3^Δ78^*) (*16, 18, 19*). This deletion results in a shift in the translational open reading frame (ORF), which results in a premature termination codon (PTC) in exon 9, upstream of both LTSs. The consequences of this premature termination event include destabilization of the mRNA due to nonsense-mediated mRNA-decay (NMD) of the mutant *CLN3* and translation of a truncated CLN3^Δ78^ protein lacking 257 amino acids at its C-terminal end, as well as both LTSs, resulting in its retention in the endoplasmic reticulum (*16, 18, 20*).

*CLN3* pre-mRNA undergoes extensive alternative splicing, with over 60 identified spliced isoforms (*21, 22*). Given the lack of knowledge about the function of the protein, it is not known which isoforms carry out the essential functions of the protein. Thus, correcting the open reading frame of *CLN3^Δ78^* and recovering the expression of at least some of the potential CLN3 protein isoforms may have therapeutic value (*18, 23, 24*). One *CLN3* spliced isoform of particular interest is *CLN3^Δ578^*which, in addition to lacking exons 7 and 8, also lacks exon 5, and is a naturally-occurring spliced isoform identified in some regions of the brain (*21, 22*). The additional skipping of exon 5 results in an intact open reading frame when exons 7 and 8 are also skipped out, resulting in a protein with a small, internal deletion, but still retaining its C-terminal end along with both LTSs. Notably, this alternatively spliced transcript has been annotated in non-human primates (ENSNLET00000044463, ENSMMUT00000001013, ENSMFAT00000017263) and dogs (ENSCAFT00845004450) in addition to humans (ENST00000357806). Conservation of this spliced isoform across species suggests it may have a functional role in cells (*23*). Having identified a naturally-occurring isoform of *CLN3* lacking exons 5, 7 and 8, we have explored strategies to restore expression from *CLN3^Δ78^* by inducing a spliced isoform with an intact open reading frame.

Splice-switching ASOs are short, synthetic nucleic acids that modulate pre-mRNA splicing by base-pairing to RNA sequences that are critical for directing appropriate splicing, thereby modulating splice-site selection (*25*). Splice-switching ASOs have been approved by regulatory agencies for the treatment of numerous genetic diseases and many more are under development. A clinical trial utilizing an intravitreal splice-switching ASO to treat the ocular disorder *CEP290*-associated Leber congenital amaurosis (LCA) demonstrated efficacy in most patients (*26, 27*). ASO clinical trials for vision loss in other diseases including Usher syndrome type 2 (*28*), and retinitis pigmentosa are being developed (*29*).

We have shown previously that intracerebroventricular (ICV) delivery of an ASO that induces *Cln3* exon 5 skipping during pre-mRNA splicing is therapeutic in a mouse model of CLN3 Batten disease, decreasing abnormal histopathology throughout the brain and improving sensorimotor coordination (*16*). We have validated these findings in a mouse model that expresses only the *Cln3^Δ578^*isoform, mimicking our ASO approach, and found that open reading frame correction by exon 5 skipping in *Cln3^Δ78^* results in a protracted disease progression in mice (*30*).

To explore whether ASO-mediated exon 5 skipping can also be used to treat retinal dysfunction in CLN3 disease, we tested the approach in a *CLN3^Δ78^* minipig model (*31*). Though mouse models have been used extensively to study this disease, rodent models of central vision loss are limited due to their lack of the highly-specialized central retina forming the macula and fovea, which are found in the eyes of humans and other primates (*32*). Furthermore, *Cln3^Δ78^*mice have limited retinal dysfunction and normal ERGs (*33, 34*). Porcine models are more appropriate for the study of retinal dystrophies given that the pig retina more closely resembles the human retina (*32, 35–39*). Indeed, *CLN3^Δ78^* pigs have progressive vision loss, retinal pigment epithelium dysfunction, and photoreceptor cell loss (*31*).

We administered ASOs targeting pig exon 5 via intravitreal injection (IVI) and found that a single treatment of exon 5-targeted ASO induced robust exon skipping for up to a year in pigs with the *CLN3^Δ78^*allele. Treatment also protects against retinal dysfunction at early stages of disease in the treated eye. To test human-targeted ASOs in the eye, we screened ASOs that were complementary to a region of *CLN3* conserved between humans and pigs and identified a pan-human/pig ASO that induces exon skipping of endogenous *CLN3* exon 5 in both patient-derived fibroblasts and in the pig eye. Taken together, our findings demonstrate the utility of ASO-based reading frame correction as an approach to treat CLN3 Batten disease retinal dysfunction and demonstrate the activity of an ASO-specific for human *CLN3^Δ78^ in vivo*.

## RESULTS

### *CLN3^Δ78^* expression in minipigs mirrors *CLN3^Δ78^* in humans

As a first step in utilizing the *CLN3^Δ78^* minipig as an *in vivo* model for ASO screening, we analyzed the sequence of the *CLN3* locus surrounding the deletion and the expression of the gene. We isolated genomic DNA (gDNA) from *CLN3^Δ78^* minipig tissue and confirmed that the model has a 952 base pair deletion between intron 6 and intron 8 of *CLN3* and a 122 base pair insertion into intron 8 (**Figure S1A,C**). The deletion begins 278 bases upstream of intron 6, encompasses the entire 297 bases of exon 7, intron 7, and exon 8, and extends 377 bases into the 3’ end of intron 8 (**Figure S1A**). Immediately following this deletion there is an insertion of 122 bases of the PGK-NeoR cassette, a vestige of the rAAV-mediated gene targeting used to generate this genetic model (**Figure S1A**). PCR analysis of gDNA from the gene-edited pig tissue revealed an amplicon of the expected size resulting from the 952 base pair deletion (**Figure S1C**). Reverse transcription-polymerase chain reaction (RT-PCR) analysis of RNA isolated from the retina and sequencing of the resulting cDNA confirmed that the deletion/insertion results in the properly-spliced *CLN3^Δ78^*mRNA transcript of the expected size (**Figure S1B,D**). These results indicate that the *CLN3^Δ78^* minipig model accurately recapitulates the most common disease-causing variant in humans with CLN3 disease.

### Identification of ASOs that induce pig *CLN3^Δ78^* exon 5 skipping *in vitro*

To identify ASOs that induce exon 5 skipping in pig *CLN3^Δ78^*(*pCLN3*), a series of twenty-four 18-mer fully-modified phosphorothioate steric blocking ASOs with 2’methoxyethyl (MOE) modifications were designed to base-pair with exon 5 and its flanking intronic region of *pCLN3* RNA (**Figure 1A**). ASOs were transfected into a fibroblast cell line derived from the skin of a homozygous *CLN3^Δ78^* minipig. Prior to collection, cells were treated with puromycin to inhibit nonsense-mediated mRNA decay to stabilize the PTC-containing *CLN3^Δ78^* mRNA and thus more accurately assess the processing of RNA expressed from the gene. RNA was analyzed by RT-PCR using primers directed to exons 4 and 6 to quantify mRNA with exon 5 skipped. Many ASOs induced robust exon 5 skipping. However, treatment with some of the ASOs appeared to result in lower levels of total CLN3 mRNA. To determine whether this lower abundance was due to the additional skipping of exon 6, we performed the PCR analysis with primers in exons 4 and 10. Surprisingly, the pig fibroblasts have a high level of exon 6 skipping in untreated cells (**Figure S2**). Skipping of exon 6 in the context of the deletion of exons 7 and 8 results in an intact open reading frame. The exon 5-targeted ASOs induced skipping of exon 5 in these transcripts as well (**Figure S2**). ASO-16 induced one of the highest levels of exon 5 skipping in each of these assessments (81% and 91%, respectively) and was selected for further study (**Figures 1B, S2**). ASO-16 base-pairs to a sequence in the middle of exon 5 and induces dose-dependent exon 5 skipping with an EC_50_ (half maximal effective concentration) in the low nanomolar range (5.39 nM) (**Figure 1C-E**). These results identify ASOs that effectively induce *pCLN3* exon 5 skipping *in vitro*.

**Figure 1.**
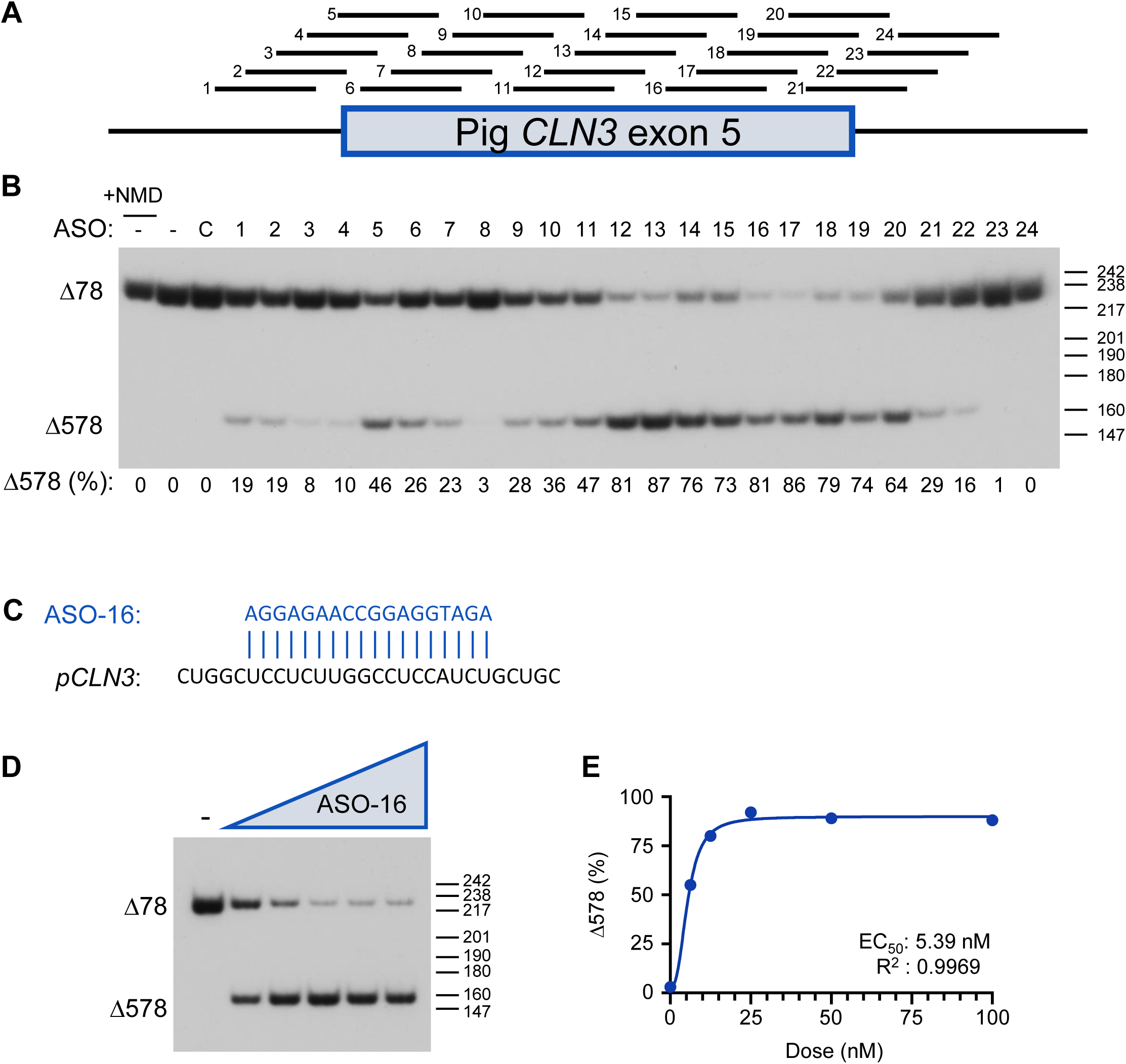
ASOs potently induce exon 5 skipping of pig *CLN3^Δ78^ in vitro*. (A) Diagram of the location of ASOs 1 to 24 on the pig *CLN3 (pCLN3)* exon 5 pre-mRNA. The box indicates exon 5 and the bars indicate the flanking introns. (B) Radioactive RT–PCR was performed on RNA extracted from *CLN3^Δ78^* cells individually transfected with the indicated ASO, mock treated (C), or untreated (–). PCR was performed using primers in *CLN3* exons 3 and 6. The spliced products are labeled on the left of the gel. Below: quantification of exon 5 splicing (calculated as a percentage *CLN3^Δ578^*/(*CLN3^Δ78^*+*CLN3^Δ578^*)) from a single experiment shown in the gel. Cells were treated with puromycin prior to collection to block nonsense mediated decay (NMD) with the exception of cells from sample labelled “+NMD”. Size markers (bp) are shown on the right of the gel. (C) The nucleotide sequence of the optimal ASO (blue) aligned with the target p*CLN3* region. (D) RT–PCR analysis of exon 5 skipping using RNA isolated from *CLN3^Δ78^*pig fibroblasts treated with increasing doses of ASO-16 (6.25 to 100 nM) or with 100 nM of a control ASO (–). Prior to collection, cells were treated with puromycin to inhibit NMD. Spliced products are labeled on the left of the gel. Size markers (bp) are shown on the right of gel. (E) Exon 5 skipped (%) relative to the dose is plotted. The half-maximal effective concentration (EC_50_) was calculated after fitting the data using non-linear regression with a variable slope.

### ASO-16 induces long-lasting exon 5 skipping throughout the retina of heterozygous pigs carrying the *CLN3^Δ78^* deletion

After identifying potent ASO-16 exon 5 skipping activity *in vitro*, we proceeded to evaluate its activity *in vivo*. One of three different doses (80, 200 or 300 μg) of ASO-16 or vehicle Dulbecco’s phosphate buffered saline (DPBS) was delivered to the retina of heterozygous *CLN3^+/Δ78^* minipigs by IVI. The 300 μg dose was selected based on previous studies that injected ASOs in the eye (*40–42*). Pigs aged 11-30 days (300 and 200 μg) or 125-154 days (80 μg dose) were treated and eye tissue and RNA from the retina were collected three, six, and twelve months later. Because there are no reliable antibodies that specifically detect CLN3 protein, our evaluation of ASO activity was limited to splicing analysis of the mRNA. RT-PCR analysis of retinal RNA revealed that at three and six months after the high dose treatment (300 μg), 98% and 91% of the total mRNA had undergone ASO-induced exon 5 skipping, respectively (**Figure 2A,B, lanes 1-4**). RNA from optic nerve was also analyzed and showed no evidence of ASO-induced exon 5 skipping (**Figure S3C**). Similar to the high dose, the intermediate dose (200 μg) dose induced 99% and 93% exon skipping at 3 and 6 months post-IVI respectively, and exon 5 skipping was maintained at 74% at 12 months post-IVI (**Figure 2A,B, lanes 5-10**). For the low dose (80 μg), 77% and 76% exon 5 skipping was observed at 3 and 6 months post-IVI, respectively, but dropped to 34% 12 months post-IVI (**Figure 2A,B, lanes 11-16**).

**Figure 2.**
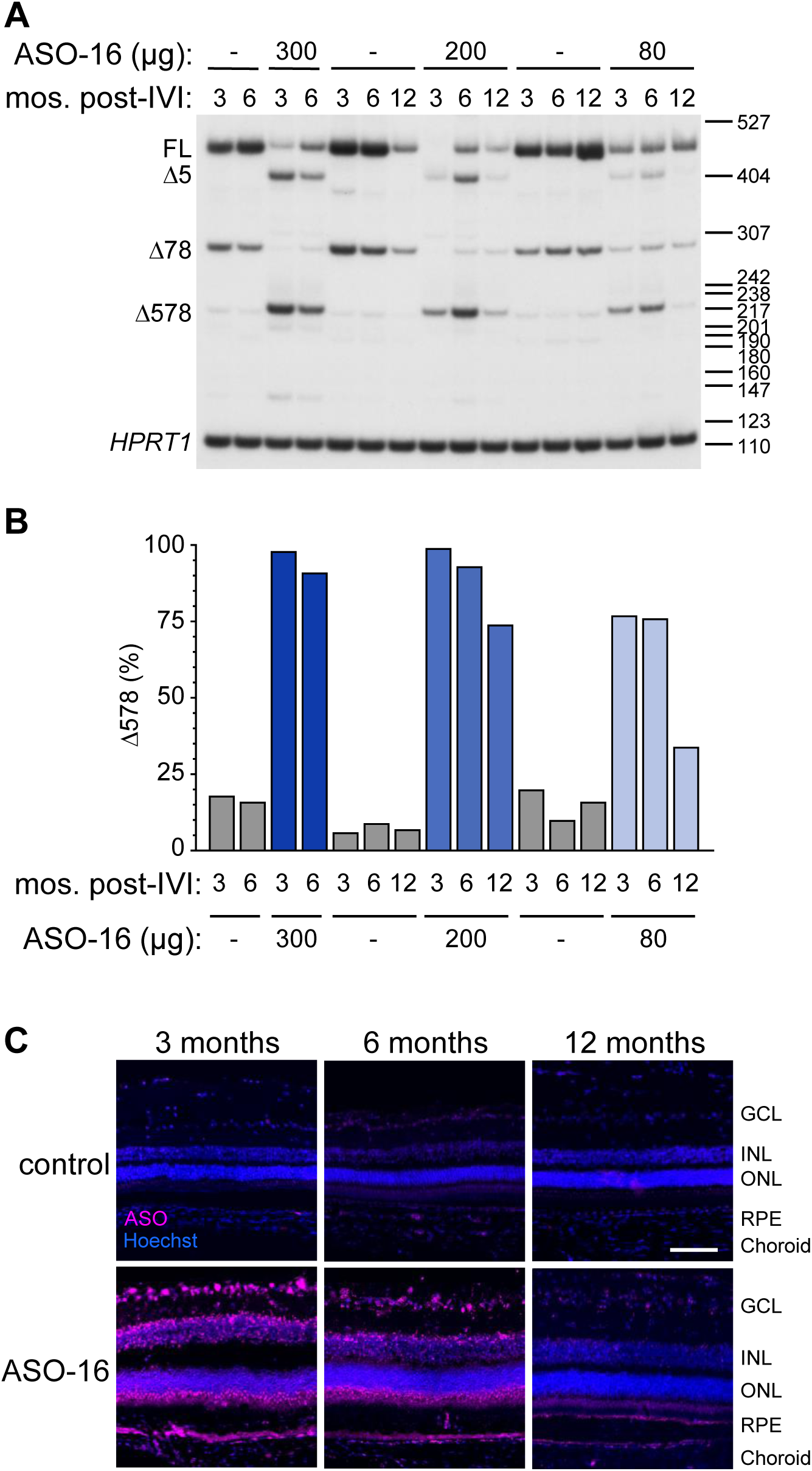
ASO-16 induces long-lasting exon 5 skipping in the retina of heterozygous *CLN3^+/Δ78^* pigs. (A) Radioactive RT–PCR analysis of exon 5 splicing in the retina of heterozygous *CLN3^+/Δ78^* pigs treated with ASO-16 or DPBS vehicle control (–) and analyzed at 3, 6 and 12 months (mos.) post-treatment. *HPRT1* was included as a loading control. Products are labeled on the left of the gel. Size markers (bp) are shown on the right of gel. (B) Quantification of exon 5 skipping in groups shown in (A). (C) Representative immunofluorescence images of ASO (magenta) and the nuclear marker Hoechst (blue) in the retina of *CLN3^+/Δ78^* pigs 3, 6, or 12 months (mos.) following a single treatment of 80 μg ASO-16 by IVI, and a contralateral, DPBS control-treated eye. Retinal layers are labeled for clarity: ganglion cell layer (GCL), inner nuclear layer (INL), outer nuclear layer (ONL), retinal pigment epithelium (RPE), and choroid. Scale bar, 100 µm.

Immunofluorescent analysis of ASO in the retina of the treated pigs detected using an ASO-specific antibody confirmed that the ASO was distributed throughout the retina, correlating with exon 5 skipping activity (**Figure 2C**). Taken together, these results reveal exon 5 skipping for up to 12 months, demonstrating the long-term efficacy of ASOs in the retina.

The treatment was generally well-tolerated with few adverse events (AEs). One vehicle-treated eye developed a corneal ulcer, one vehicle-treated eye developed a cloudy vitreous, and one vehicle-treated eye developed a cataract (**Table S1**). Though these were heterozygous pigs and not expected to have any visual deficits associated with loss of one copy of *CLN3*, ERGs were performed on the pigs to evaluate any adverse effects of the treatment on retinal function. ERGs of eyes treated with the low dose of ASO (80 μg) had measurements similar to vehicle-treated eyes, whereas eyes treated with the higher doses had diminished ERG amplitudes following treatment, which recovered over time (**Table S1**). Diminished amplitudes could be caused by a number of factors including the dose of the ASO itself, the younger age of the pigs treated with the higher doses at the time of treatment (11-33 days old vs 125-154 days old for the 80 μg dose), or the knock-down of the wild-type CLN3 allele in these heterozygous pigs caused by ASO-induced exon 5 skipping. To address these possibilities and limit the risk of adverse events while still treating early enough to stabilize *CLN3* transcript expression when it is highest (**Figure S3**), we moved forward with a conservative treatment protocol to test ASO efficacy, administering the low dose of ASO (80 μg) no earlier than 3.4 months of age.

### A single intravitreal injection of ASO protects against retinal dysfunction in homozygous *CLN3^Δ78^* pigs

To test the effect of ASO treatment in homozygous *CLN3^Δ78^*pigs, we performed a longitudinal study to determine whether ASO-16 can induce exon 5 skipping and prevent retinal dysfunction. ASO-16 (80 μg) was delivered to the retina of homozygous *CLN3^Δ78^* minipigs via a single IVI, with the contralateral eye serving as a vehicle-treated control. Heterozygous *CLN3^+/Δ78^* pigs were treated in one eye with vehicle by IVI to control for injection-related effects and measure unaffected vision over time. All pigs were treated between 3.4 and 3.9 months of age. ASO-16 was well-tolerated in the retina (**Figure S4**). None of the animals treated with ASO-16 had ocular AEs, though one control animal developed a partial cataract in its vehicle-treated eye (**Table S1**).

To evaluate whether ASO-16 could prevent retinal dysfunction in *CLN3^Δ78^*minipigs, we performed ERGs to measure the electrical response of the retina to light stimulation. ERGs were performed immediately before injection and every 3 months post-treatment for 9 months (**Figure 3A**). Testing was performed under light-adapted and dark-adapted conditions to assess the cone-predominant and combined cone and rod response, respectively. Photostimulation with 8.0 cd•s/m^2^ (bright flash) and 25 cd•s/m^2^ (super-bright flash) was used and the resulting photoreceptor predominant (a-wave) and bipolar cell predominant (b-wave) responses were measured (**Figure 3B-D**). The implicit time of the a-wave and b-wave, which represents the speed of response of the cells following stimulation (*43*), was not significantly different between ASO-treated vs vehicle-treated eyes at any timepoint or testing condition (**Figure S5**).

**Figure 3.**
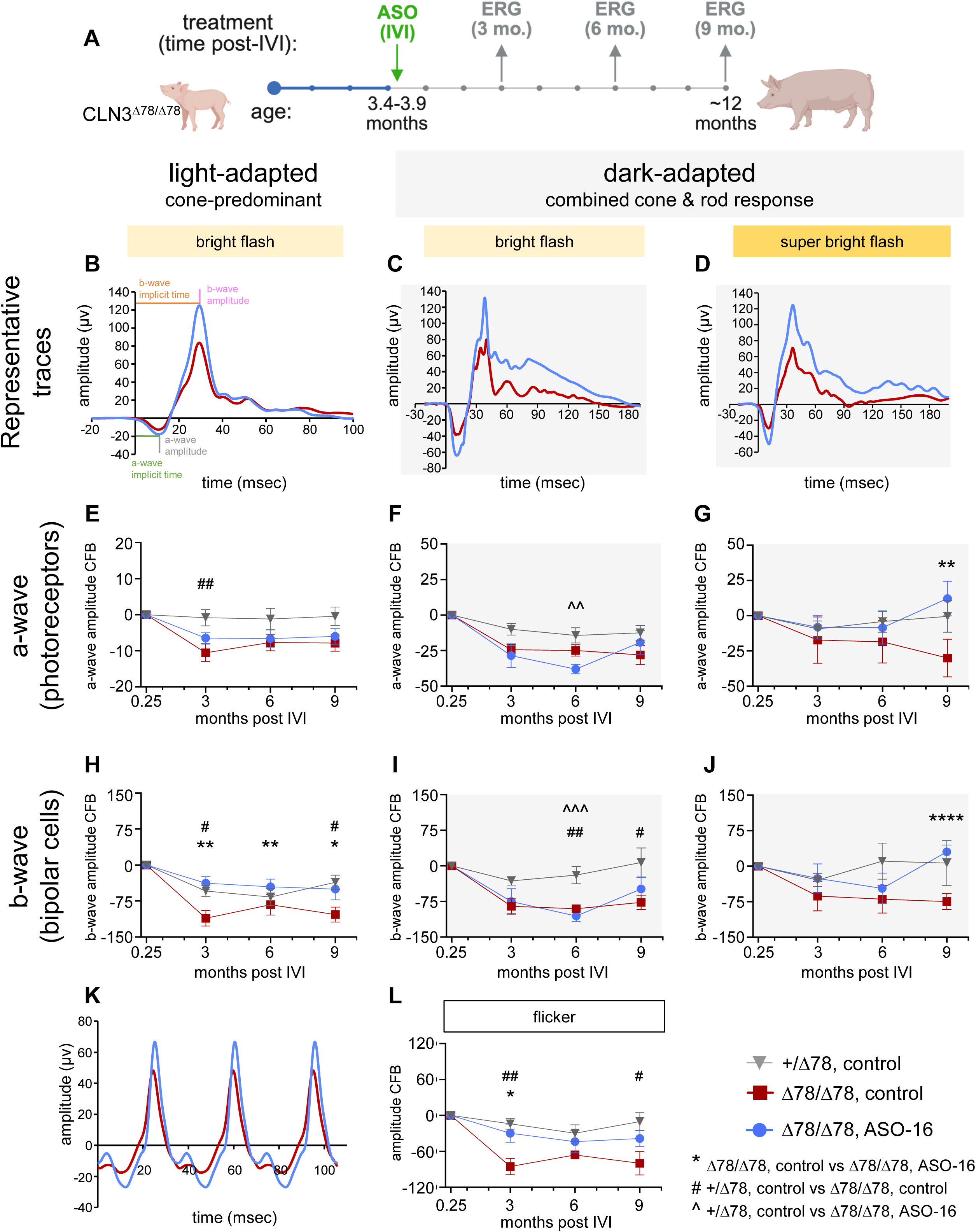
A single intravitreal injection of ASO-16 protects against retinal dysfunction in *CLN3^Δ78^* pigs. (A) Schematic showing the timeline of ASO treatment and ERG analyses. Image created with www.biorender.com. (B-D) Representative ERG traces comparing the ASO-treated (ASO-16) and DPBS vehicle-treated, control eye of *CLN3^Δ78^* pigs 9 months post-treatment. The change from baseline (CFB) in ERG (E-G) a-wave and (H-J) b-wave, comparing vehicle-treated control eye of *CLN3^+/Δ78^* and *CLN3^Δ78^* pigs and 80 µg ASO-16 treated *CLN3^Δ78^* pigs over time with bright flash (8.0 cd•s/m^2^) or super bright flash (25.0 cd•s/m^2^). (K) Representative ERG traces comparing the ASO-treated and vehicle-treated control eye of *CLN3^Δ78^* minipigs 9 months post-treatment. (L) The change from baseline (CFB) in ERG amplitudes at 28.3 Hz. Bars show s.e.m.; one-tailed paired t-test comparisons at each timepoint of vehicle treated *CLN3^Δ78/Δ78^* and 80 µg ASO-16 \**P* <0.05, \*\**P* <0.01, \*\*\*\**P* <0.0001; one-way ANOVA with Dunnett’s multiple comparisons at each time-point of vehicle-treated *CLN3^+/Δ78^* to vehicle-treated *CLN3^Δ78^* ^(#^P<0.05, ^##^P<0.01) and 80 µg ASO-16 treated *CLN3^Δ78^* (^^P<0.01, ^^^P<0.001).

The ERG a-wave hyperpolarization “negative peak” of an ERG waveform reflects the function of the photoreceptors in the outer retina (*43*). Under light-adapted conditions with an 8.0 cd•s/m^2^ flash, the only timepoint at which a difference was observed in the cone-predominant photoreceptor system between the groups was at 3 months post-treatment when the vehicle-treated *CLN3^Δ78^* eyes were significantly worse than heterozygous control eyes (**Figure 3E, Table S2**). Although the photoreceptors of the ASO-treated *CLN3^Δ78^* eyes functioned similarly to the heterozygous control eyes at this same timepoint, the improvement compared to their contralateral, vehicle-treated *CLN3^Δ78^* eyes was not statistically significant (p = 0.0570). Under dark-adapted conditions with an 8.0 cd•s/m^2^ bright flash, the combined cone and rod photoreceptor response was not significantly different among any of the groups over the duration of the study, except for a significant drop in a-wave amplitude in the ASO-treated eyes 6-months post-treatment (**Figure 3F, Table S2**). However, at 9 months post-treatment the amplitudes from eyes treated with ASO were not different from heterozygote control eyes under these conditions (**Figure 3C, F, Table S2**). Under similar dark-adapted conditions, but with a brighter flash (25.0 cd•s/m^2^) to recruit more cone photoreceptors, the a-wave response of the ASO-treated eyes was improved significantly compared to the vehicle-treated eyes of the *CLN3^Δ78^*pigs at 9 months post-treatment (**Figure 3D,G, Table S2**). Taken together, these results indicate that a-wave amplitudes may not be profoundly reduced in *CLN3^Δ78^* pigs at this stage of disease progression, but ASO-16 treatment prevented progressive photoreceptor dysfunction, as evidenced by the significant difference in amplitudes compared to controls under super bright light stimulus at 9 months post-treatment.

The ERG b-wave depolarization “peak” reflects the function of the bipolar cells and Müller cells of the inner layers of the retina (*43*). Under light-adapted conditions with an 8.0 cd•s/m^2^ flash, cone-predominant bipolar cell function (b-wave) of vehicle-treated *CLN3^Δ78^* eyes was significantly diminished compared to healthy controls at 3 and 9 months post-treatment.

Importantly, b-wave amplitudes were significantly higher in homozygote *CLN3^Δ78^* eyes treated with ASO-16 compared to vehicle-treated eyes at all timepoints (**Figure 3H, Table S2**). Under dark-adapted conditions with an 8.0 cd•s/m^2^ bright flash, the combined cone and rod photoreceptor responses were not significantly different between the ASO-treated and vehicle-treated *CLN3^Δ78^* eyes over the duration of the study (**Figure 3C,I, Table S2**). Both the ASO– and vehicle-treated eyes of the homozygous *CLN3^Δ78^* pigs were significantly lower than the heterozygote controls in b-wave amplitude at 6-months post-treatment. However, at 9 months post-treatment, the b-wave amplitudes of ASO-16-treated eyes from homozygote *CLN3^Δ78^*pigs were not significantly different than heterozygote control eyes, and the vehicle-treated *CLN3^Δ78^* eyes were significantly lower (**Figure 3I, Table S2**). Similar to the a-wave results, the b-wave response with the 25.0 cd•s/m^2^ super bright dark-adapted flash was significantly higher in the *CLN3^Δ78^* homozygote ASO-treated eyes compared to the vehicle-treated eyes at 9 months post-treatment (**Figure 3D,J, Table S2**).

Using the light-adapted 28.3 Hz flicker ERG, the bipolar cells of the cone-only pathway functioned significantly better in the ASO-treated eye at 3 months post-treatment, though this difference was not significant at 9 months post-treatment (**Figure 3K,L, Table S2**). In contrast, the vehicle-treated eye of the *CLN3^Δ78^* homozygote pigs were significantly decreased compared to healthy controls at both 3 and 9 months post-injection (**Figure 3L, Table S2**). Taken together, these results suggest that b-wave amplitudes may be more affected than a-wave amplitudes at this stage of disease in pigs, and that ASO-16 may stabilize function in both cone and combined photoreceptor responses, particularly under light-adapted and super bright flash conditions.

### Improved retinal function in a subset of high-responding animals treated intravitreally with ASO-16

Among the seven *CLN3^Δ78^*pigs that were treated with ASO-16 and were included in the final study, a subset of four animals were identified as high responders, a phenomenon seen in human studies with ASOs following IVI injection (*44*). Animals were classified as high-responders if they showed improvement in their ASO-treated eye in at least six of the seven ERG measurements recorded at 9 months post-treatment (**Figures 4, S6, S8, Tables S3,S4**). In these animals, under dark-adapted conditions, the combined rod and cone photoreceptor response was significantly improved using both a 8.0 cd•s/m^2^ bright flash and 25.0 cd•s/m^2^ super bright flash (**Figures 4B,C, S6C,D, Table S3**). Bipolar cell function was also significantly improved in the cone-only (**Figure 4G, S6A, Table S3**), cone-predominant (**Figure 4D, S6B, Table S3, Table S3**), and combined cone and rod pathway (**Figure 4F, S6D, Table S3**) of this subgroup, although improvement in the combined pathway was only seen with the 25.0 cd•s/m^2^ super bright flash (**Figure 4,E,F, S6D, Table S3**). Taken together, these results indicate that some animals responded more favorably to *CLN3* open reading frame correction than others.

**Figure 4.**
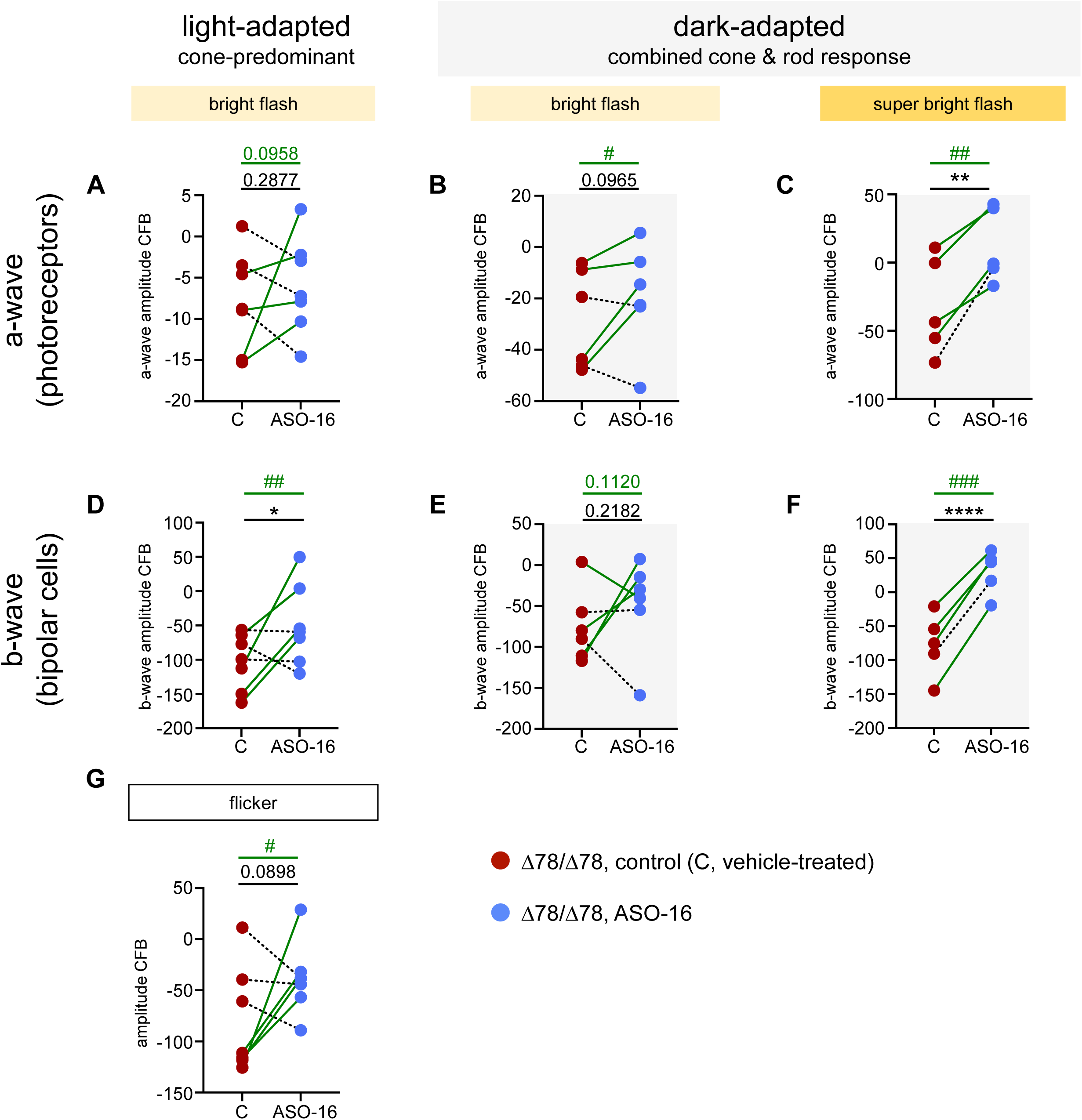
Preserved retinal function in a subset of high-responding animals treated intravitreally with ASO-16. (A-G) The change from baseline (CFB) in ERG amplitudes of individual *CLN3^Δ78^* pigs 9 months post-treatment (A-C) a-wave, (D-F) b-wave, and (G) amplitudes at 28.3 Hz flicker. Comparing vehicle-treated, control-treated and ASO-16 treated (80 µg) eyes of each *CLN3^Δ78^*pig with bright flash (8.0 cd•s/m^2^) or super bright flash (25.0 cd•s/m^2^). Connecting lines indicate data points corresponding to ASO-16 and control, contralateral eye of the same animal, with green lines highlighting high-responding animals. One-tailed paired t-test comparisons for all animals shown in black; \**P* <0.05, \*\**P* <0.01, \*\*\*\**P* <0.0001; one-tailed paired t-test comparisons for high-responding animals shown in green; ^#^P<0.05, ^##^P<0.01, ^###^P<0.001.

A subset of the cohort of pigs in the longitudinal study (homozygous CLN3^Δ78^ only) were maintained until 12 months post-injection and ERGs were measured (**Figure S9, Table S5**).

There was no significant difference between control and ASO-16-treated eyes, except in the high-responders identified in the longitudinal study (Figure 4), whose ASO-treated eyes were significantly improved in the flicker response (**Figure S9G**) To confirm that ASO-16 induced exon 5 skipping in the homozygote *CLN3^Δ78^* pig eyes, we analyzed RNA from three of the pigs at 12 months post-injection. The remainder of the pigs were maintained for ongoing, long-term functional testing and re-dosing experiments. RT-PCR analysis of retinal RNA isolated from the pigs confirmed exon 5 skipping (54%), verifying the delivery and long-term durability of ASOs following IVI (**Figure S9H,I**). Overall, we conclude from these results that the positive effect of ASO treatment on ERGs was largely limited to 9 months post treatment, with minimal benefit, relative to the vehicle-treated eye at 12 months post-treatment (**Figure S9**). The decline at 12 months post-injection may be due to the waning effect of the ASO on splicing from 6 to 12 months post-treatment, a decrease that we extrapolate from the experiments with heterozygote pigs, wherein ASO activity was steady up until six months post-injection but dropped dramatically at 12 months post-injection (**Figure 2**).

### Splice-switching ASOs targeting a region of *CLN3* conserved across pigs and humans induce exon 5 skipping in human *CLN3^Δ78^* fibroblasts

Having demonstrated the ability of an ASO to induce *CLN3^Δ78^* exon 5 skipping in the retina and mitigate retinal dysfunction in homozygote *CLN3^Δ78^* pigs, we sought to extend these findings to a clinically-relevant ASO that has the potential to target human RNA and can be tested in the pig model of the disease to demonstrate the efficacy of a possible drug candidate. In addition to recapitulating the loss of ERG amplitude phenotype seen in patients with CLN3 Batten disease, the *CLN3^Δ78^*minipig model has the advantage of increased sequence homology with the human gene. Indeed, a region encompassing the 5’ splice site of exon 5 is highly homologous between humans and pigs.

To identify an active human *CLN3*-targeted ASO within the region homologous between humans and pigs, we tested four conserved 18-mer ASOs (**Figure 5A,C**). The ASOs were transfected into a homozygous *CLN3^Δ78^* patient-derived fibroblast cell line and splicing was analyzed by RT-PCR. All four ASOs induced nearly 100% exon 5 skipping (**Figure 5B**).

**Figure 5.**
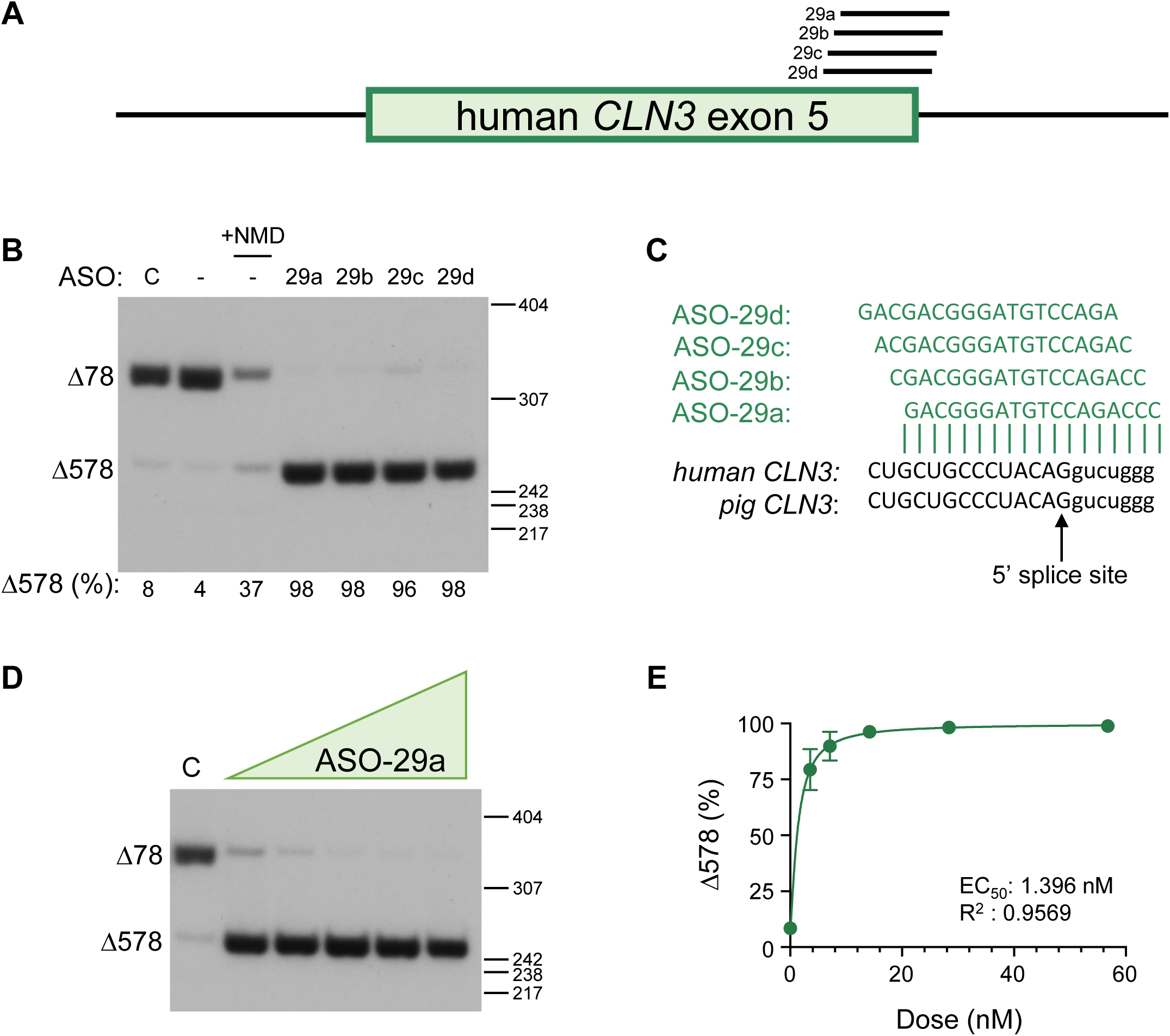
Splice-switching ASOs targeting a region of *CLN3* conserved across pigs and humans induce exon 5 skipping in human *CLN3^Δ78^* fibroblasts. (A) Diagram of the location of ASOs 29a to 29d on the human *CLN3 (hCLN3)* exon 5 pre-mRNA. The box indicates exon 5 and the bars indicate the flanking introns. (B) Radioactive RT–PCR was performed on RNA extracted from *CLN3^Δ78^* cells individually transfected with 100 nM of the indicated splice-switching ASO. The spliced products are labeled on the left of the gel. Below: quantification of exon 5 splicing (as a percentage of total *CLN3^Δ78^* mRNA). Control-non-targeted, ASO-treated (C) and untreated (–) controls are included. Cells were treated with puromycin prior to collection to inhibit NMD, unless indicated (+NMD). Size markers (bp) are shown on the right of gel images. (C) The nucleotide sequence of the ASOs aligned with the target *pCLN3* and *hCLN3* region with exon 5 sequence in capital letter and intron 5 sequence in lowercase letters. (D) RT– PCR analysis of exon 5 skipping using RNA isolated from *CLN3^Δ78^* cells treated with a control ASO (C) or increasing doses of ASO-29a (3.5 to 57 nM) and 5-6 h prior to collection, treated with puromycin. Products are labeled on the left of the gels. Size markers (bp) are shown on the right of gels. (E) Exon 5 skipped (%) in relationship to the dose is plotted from 3 independent experiments. The half-maximal effective concentration (EC_50_) was calculated after fitting the data using non-linear regression with a variable slope.

To select an ASO for testing *in vivo*, we performed a limited tolerability test in rats by IVI of each ASO. We selected ASO-29a for further study based on its favorable tolerability profile, though all tested ASOs were well-tolerated (**Figure S10**). ASO-29a base pairs across the 5’ splice site of exon 5 and has an EC_50_ *in vitro* in the low nanomolar range (1.396 nM) (**Figure 5D,E**). These results demonstrate the ability of ASOs to target a human *CLN3* (*hCLN3*) sequence and induce robust exon 5 skipping in the *CLN3^Δ78^* minipig model.

### ASO-29a induces stable, long-lasting *CLN3* exon 5 skipping throughout the retina

To assess the *in vivo* activity of ASO-29a, homozygous *CLN3^Δ78^* minipigs were treated between 103 and 178 days of age with a single IVI of ASO-29a (80 μg or 160 μg) or a vehicle control.

ERGs were taken one-week, three-months, and six-months post-treatment to evaluate safety and tolerability of the treatment (**Figure S11**). Tissue and RNA were collected to assess ASO distribution and exon skipping. One eye treated with 160 μg of ASO-29a and one eye injected with vehicle developed cataracts (**Table S1**). With the exception of a decrease in the 30 Hz flicker ERG response 3 months post-treatment, ASO-29a was well-tolerated in the retina at 80 μg, but was associated with diminished ERGs at the 160 μg dose (**Figure S11**). For the 80 μg dose, ASO-29a induced exon 5 skipping in 87%, 95%, and 95% of *CLN3* RNA transcripts, one, three, and six months after treatment respectively (**Figure 6A,B, lanes 6-10**). Similarly, the 160 μg dose resulted in 99% exon 5 skipping at all three timepoints (**Figure 6A,B, lanes 16-20**).

**Figure 6.**
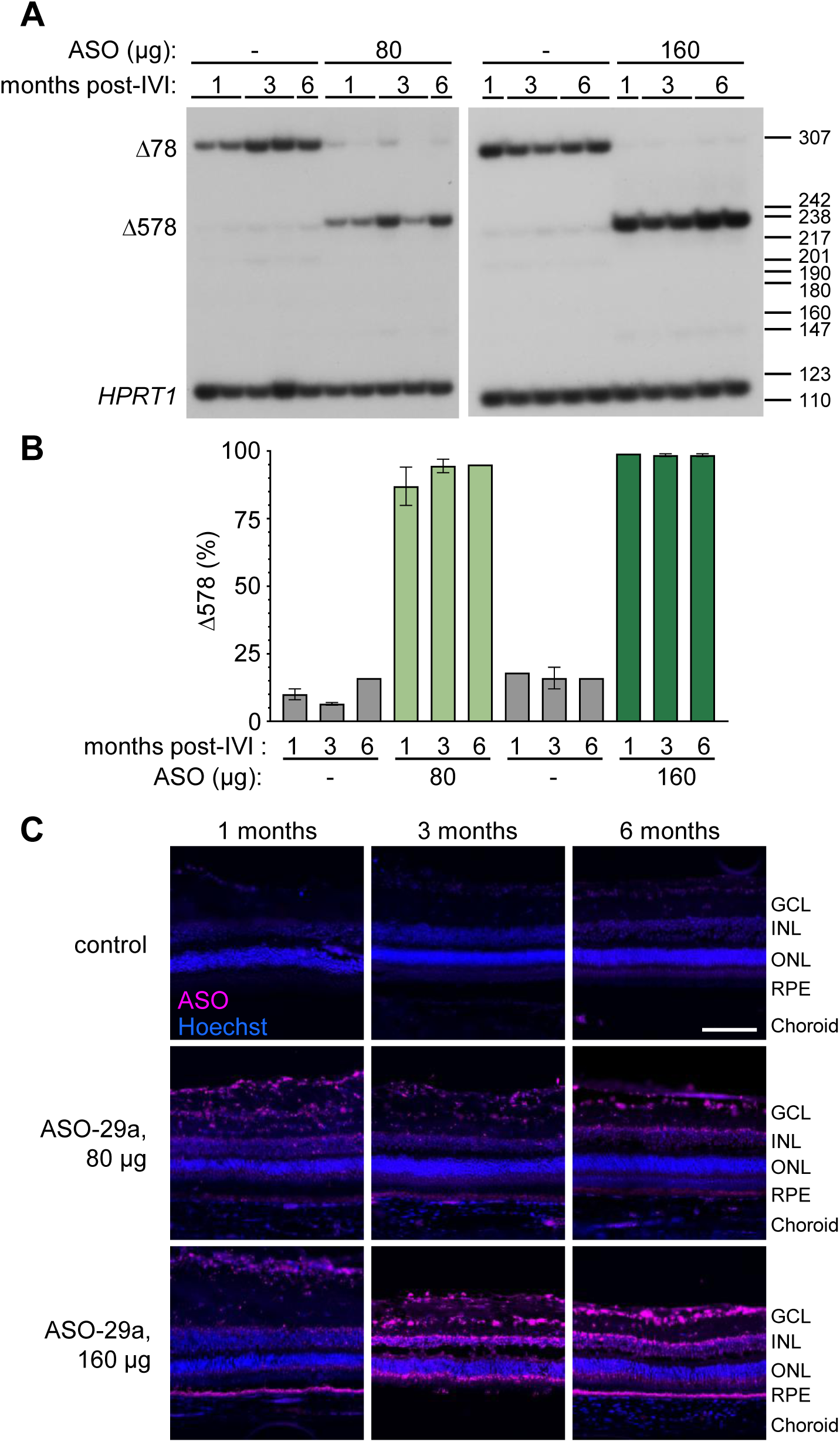
ASO-29a induces stable, long-lasting *CLN3* exon 5 skipping throughout the retina. (A) Radioactive RT–PCR analysis of exon 5 splicing in the retina of homozygous *CLN3^Δ78^* pigs treated with ASO-29a or vehicle (–) and analyzed at 1 month, 3 months, and 6 months post-treatment. *HPRT1* was included as a loading control. Products are labeled on the left of the gel. Size markers (bp) are shown on the right of gel. (B) Quantification of exon 5 skipping in groups shown in (A). (C) Representative immunofluorescence images of ASO (magenta) and the nuclear marker Hoechst (blue) in the retina of *CLN3^Δ78^*pigs 1, 3, or 6 months following a single treatment of 80 μg or 160 μg ASO-29a by IVI, and a contralateral, DPBS control treated eye. Retinal layers are labeled for clarity: ganglion cell layer (GCL), inner nuclear layer (INL), outer nuclear layer (ONL), retinal pigment epithelium (RPE), and choroid. Scale bar, 100 µm.

ASO was distributed throughout the retina and apparent abundance of the ASO correlated with exon 5 skipping activity over time (**Figure 6C**). Similar to ASO-16, ASO-29a induces robust, long-lasting exon 5 skipping for up to 6 months in the retina, demonstrating the activity of a potential clinically-relevant ASO for treating CLN3 Batten disease vision loss in humans.

## DISCUSSION

Here we demonstrate that ASO-induced skipping of *CLN3^Δ78^* exon 5 protects the retina from early functional losses in a large animal model of the most common cause of CLN3 Batten disease. We demonstrate that this therapeutic approach is well-tolerated in the eye and that a single intravitreal injection of ASO induces robust exon skipping for up to a year. Furthermore, we identify an ASO that base-pairs with 100% complementarity to both pig and human *CLN3* and is highly potent at inducing exon 5 skipping in the retina. This result is an important demonstration of *in vivo* target engagement of an ASO that could be considered for treating CLN3 disease in humans.

For a therapeutic to be effective, it is essential that the treatment is active in the cells that are affected in the disease. In patients with CLN3 Batten disease, there is severe loss of the photoreceptor ONL layer in the macula and mid-periphery (*10, 45, 46*), along with the reduced ERG a-wave responses (*9–13*), suggesting that photoreceptor cells are profoundly affected by CLN3 disease. The effect of CLN3 loss on photoreceptors could be direct or indirect through the dysfunction of retinal pigment epithelium cell phagocytosis of the photoreceptor outer segment (*47*). Additionally, bipolar cell rescue may play an important role in rescuing retinal degeneration given that patients exhibit even greater loss of b-wave amplitudes (*11*) and AAV therapy targeting bipolar cells, but not photoreceptors, was found to be therapeutic in *Cln3^Δ78^* mice (*48*).

Our results indicate that ASO-16 reached both the inner and outer retina (**Figures 2C, 6C**), and while photoreceptor and bipolar cell function both showed improvement with treatment, this effect was more pronounced in bipolar cells (**Figure 3**). Given the implication of both photoreceptors and bipolar cells in CLN3 disease vision loss, further investigation is warranted into how ASO treatment impacts function and survival of both of these cell types. Furthermore, the greatest improvements, both in photoreceptors and bipolar cells, were seen with the cone-predominant light-adapted ERGs or brighter 25.0 cd•s/m^2^ dark-adapted flash, which recruits more cone photoreceptors. These improvements, which were largely absent from the rod-predominant 8.0 cd•s/m^2^ dark-adapted flash, could indicate that ASO-16 has a greater therapeutic benefit in bipolar cells and cone photoreceptors vs rod photoreceptors. However, given that these pigs were early in their disease progression, it is possible that it was too early to see degeneration in the rod pathway, whereas we observed early-stage degeneration in the cone pathway, which was prevented through ASO treatment. A more long-term study in these animals could reveal whether ASO treatment stabilizes rod pathway function in a similar manner.

The eye is an attractive organ for translating therapeutics from the lab to the clinic for many reasons (*49*). The retina is a readily accessible portion of the CNS, meaning that tests of retinal function have potential value in assessing the efficacy of CNS-targeting therapies (*12*). Given the significant impact of preserving vision in just one eye on a patient’s quality of life, using one eye as an internal control for the other treated eye may be an effective strategy for clinical trials. This approach not only allows for direct comparison within each patient, but also eliminates the need to ask parents to potentially enroll their child in a placebo group. The eye also offers increased safety for patients during early clinical trials given that the small size of the eye means only small doses are needed, local delivery enhances drug bioavailability, and the eye is a closed compartment, separated from the rest of the body by the blood-retinal barrier, ensuring very little drug will circulate systemically (*49–51*).

IVIs are the most common intraocular procedure worldwide, and are standardly performed by an ophthalmologist (*52*). In addition to being routine care for the treatment of neovascular disorders of the retina, IVI has been shown to be a safe and effective route of delivery for ASOs to the retina (*26, 27, 53*). The most commonly noted side effects for IVI are short-term increases to intraocular pressure and the development of cataracts (*27, 52, 54*). While the possibility of adverse events is always of concern for patient well-being, it is worth noting that cataracts can be surgically corrected if they occur. Given the longevity that ASOs have shown intravitreally both here and by others, treatment windows of a single injection every 6-12 months should reduce the risk of adverse events while providing adequate therapeutic coverage (*55*).

The therapeutic effect of the ASO treatment 9 months after a single treatment was largely lost by 12 months post-treatment when exon 5 skipping had dropped to 54% (**Figure S9H,I**). This result is consistent with our previous findings in a mouse model that endogenously expresses the *Cln3^Δ578^*isoform, where homozygous *Cln3^Δ578^* mice had lower disease burden compared to heterozygous *Cln3^Δ78/Δ578^* mice. Together, these studies suggest that the threshold for therapeutic efficacy may occur with ≥50% exon 5 skipping (*30*).

Early therapeutic intervention is crucial for maximizing the outcomes of a progressive neurodegenerative disease with childhood onset. Nonetheless, though earlier treatment with ASOs showed superior benefit in conditions like SMA, benefit is still observed after symptom onset (*56, 57*). The greatest therapeutic effect may occur early in development when *CLN3* expression is highest in the brain (*58*) and retina (**Figure S3**). However, administering treatment too early in a developing system, such as the eye, could be harmful. Indeed, studies in neonatal rats have shown that IVI administration of ASOs can be detrimental if given too early, but beneficial if delayed by a few weeks (*59*). These findings highlight the importance of determining the optimal timing and dosage for treating children with CLN3 disease. Given similarities in retinal development between humans and pigs, minipigs are an ideal model for investigating this (*60–63*). Future studies should focus on refining treatment timing and dosage to maximize therapeutic outcomes.

For families with children who have CLN3 disease, diagnosis often does not take place until after symptoms have begun, so it is typically not a matter of how early treatment can begin but rather, how late. The current study only assessed the ability of ASO-mediated ORF correction to prevent retinal dysfunction – not reverse it. It remains to be determined whether later treatment, after retinal deficits appear, will be effective. Interestingly, a recent study utilizing an ASO to treat *CEP290*-associated LCA demonstrated improved visual acuity in a subset of patients who received the treatment after the onset of retinal dysfunction (*27*). Importantly, this same group also saw a subset of high-responders within their patient population, similar to what we have reported here in pigs (*27, 55*).

Our study demonstrates that ASO-induced exon 5 skipping of *CLN3^Δ78^*can protect against retinal dysfunction early in CLN3 disease. However, a therapy targeting long term CLN3 disease-associated vision loss may require dual delivery both intravitreally and intrathecally to correct *CLN3* expression in both the retina and visual cortex. Recent work in an ovine model of CLN5 Batten disease found that gene therapy delivered intravitreally was able to protect against retinal dysfunction, but was not sufficient to prevent vision loss, likely due to cortical blindness (*64*). However, dual delivery to the CNS and eye was able to prevent vision loss (*65*).

Our study has several limitations. Most notably, due to the cost of a longitudinal study in a large animal model, the cohort sizes in some cases are small. Specifically, the splicing longevity of ASO-16 (**Figure 2**) was assessed in a single animal for each dose and timepoint. Additionally, this study was concluded 12 months post-treatment, at a relatively early stage of retinal degeneration. A more long-term analysis will provide insight into both the optimal timing for reinjection intervals as well as therapeutic efficacy as vision continues to decline in older, untreated animals. Future investigations to explore the effects of treatment following the onset of retinal dysfunction will also be important.

Taken together, our results demonstrate the potential of ASOs as a therapeutic for treating the eye in CLN3 Batten disease. The ASOs we tested are effective in modulating *CLN3* splicing in the retina to protect against retinal dysfunction when administered pre-symptomatically and demonstrate the activity of an ASO specific for human *CLN3^Δ78^ in vivo*. Our findings highlight the therapeutic value of reading-frame correction in the eye and support further exploration of this approach for treating CLN3 Batten disease vision loss.

## MATERIALS AND METHODS

### Animals

All pigs were maintained at Exemplar Genetics (Sioux Center, IA, USA) under approved institutional animal care and use committee (IACUC) protocols (MRP 2021-014 and U Iowa 4072610). Animals were group housed, and all males included in this study were castrated. All rats were maintained at Ionis Pharmaceuticals (Carlsbad, CA, USA) under an approved IACUC protocol (2021–1164).

### DNA isolation and analysis

Genomic DNA (gDNA) was isolated from the optic nerve tissue of wild-type and *CLN3^Δ78^* minipigs using the REDExtract-N-Amp Tissue PCR Kit (Sigma-Aldrich). gDNA was PCR amplified using primers specific for pig *CLN3* (*pCLN3*) intron 6 forward and intron 8 reverse (**Table S6**). PCR products were separated on a 2% agarose gel, visualized with ethidium bromide, purified using the GFX Gel Band Purification Kit (Cytiva, Marlborough, MA), and sequenced to confirm identity (Azenta, Burlington, MA). Sequences were aligned using MultAlin multiple sequence alignment (*66*).

### RNA isolation and analysis

Optic nerve and retina tissue from wild-type, *CLN3^+/Δ78^*, and *CLN3^Δ78^*minipigs was homogenized in TRIzol reagent (Life Technologies, Carlsbad, CA) and RNA was isolated according to the manufacturer’s protocol, and subsequently reverse transcribed using GoScript reverse transcriptase (Promega, Madison, WI) with oligo-dT primers. Radiolabeled PCR of the cDNA was performed using GoTaq Green (Promega, Madison, WI) with α-^32^P-deoxycytidine triphosphate, and primers specific for pig *CLN3* exon 3 forward and exon 6 reverse or exon 4 forward and exon 10 reverse, as indicated (**Table S6**). PCR products were separated on a 6% nondenaturing polyacrylamide gel and quantified on a Typhoon phosphorimager (GE Healthcare, Chicago, IL).

### Antisense oligonucleotides

Synthesis and purification of all ASOs was performed as previously described (*67–69*). ASOs were uniformly modified with 2’-O-methoxyethyl sugars with a phosphorothioate backbone. Lyophilized ASOs were dissolved in sterile Dulbecco’s phosphate buffered saline (DPBS) without calcium chloride or magnesium chloride and sterilized through a 0.2 μm filter. The ASOs were diluted to the desired concentration required for dosing animals in sterile 1x DPBS. ASO sequences are provided in **Table S6**. BLAST and GGGenome searches with the ASO-16 target sequence against the pig genome and ASO-29a target sequence against the human genome revealed no other perfect sequence matches. Potential off-target sequences with greater than 15 contiguous bases of homology are provided in **Table S7**.

### Cell culture and transfection

Pig fibroblast cell lines were grown in Dulbecco’s Modified Eagle’s medium (DMEM) supplemented with 10% FBS in a 37°C incubator with 5% carbon dioxide. Primary homozygous *CLN3 ^Δ78^* pig cells were propagated in culture following isolation from a skin biopsy taken from homozygous *CLN3 ^Δ78^*miniswine ears. Samples were cleaned by washing with Hank’s Buffered Salt Solution (HBSS) in a 10-cm tissue culture dish. After washing, samples were cut into smaller pieces and placed in a T25 tissue culture flask containing digestions media (Dulbecco’s Modified Eagle Medium with High Glucose, 4.0 mM L-Glutamine, and Sodium Pyruvate supplemented with 20% Fetal Bovine Serum, 1.2% Penicillin-Streptomycin, 0.12% Deoxyribonuclease I, 1.2% Amphotericin B, and 1.2% Collagenase). Samples were incubated overnight (37°C, 5% CO_2_). After incubation, the supernatant was centrifuged. The pellet containing cells was then plated in a T75 tissue culture flask with culture media (Dulbecco’s Modified Eagle Medium with High Glucose, 4.0 mM L-Glutamine, and Sodium Pyruvate supplemented with 20% Fetal Bovine Serum, 1% Penicillin-Streptomycin, and 1% Amphotericin B). Once the cells started to grow, the culture media was changed to Dulbecco’s Modified Eagle Medium with High Glucose, 4.0 mM L-Glutamine, and Sodium Pyruvate supplemented with 10% Fetal Bovine Serum, 1% Penicillin-Streptomycin, and 1% Amphotericin B. When cells were ready to be expanded, 0.25% Trypsin was used to dissociate the cells from dishes with a five-minute incubation at room temperature. Trypsin was inhibited by adding Dulbecco’s Modified Eagle Medium with High Glucose, 4.0 mM L-Glutamine, and Sodium Pyruvate supplemented with 10% Fetal Bovine Serum and 1% Penicillin-Streptomycin. After centrifugation, the cells were expanded 1:2. Adherent cells were expanded for 10 passages prior to transfection.

Fibroblast cell cultures, derived from a person with CLN3 Batten disease who is homozygous for *CLN3^Δ78^* (SP3.2.1) were provided by F. Porter and A. D. Do (National Institutes of Health (NIH)/National Institute of Child Health and Human Development (NICHD)). Human fibroblasts were grown in DMEM/high glucose medium supplemented with 10% FBS.

ASOs targeting exon 5 of pig *CLN3* or human *CLN3* were individually transfected into a primary cell line from homozygous *CLN3^Δ78^*miniswine ears or fibroblasts from a patient with CLN3 Batten disease, at a final concentration of 100 nM, or for the dose response from 0 nM to 100 nM for pig cells and 0 nM to 57 nM for human cells. ASOs were transfected into cells using Lipofectamine 3000 Reagent (Thermo Fisher Scientific) according to the manufacturer’s protocol, with the P3000 reagent included only for the ASO walk in pig fibroblasts. Cells were collected in TRIzol (Life Technologies) after 48 h and analyzed for exon skipping. RNA was collected using TRIzol and analyzed by RT-PCR as described above, using primers spanning exon 3 forward and exon 6 reverse, or exon 4 forward and exon 10 reverse for pig *CLN3*; and exon 4 forward and exon 10 reverse for human *CLN3*. Where indicated, cells were treated with puromycin at a final concentration of 200 µg/mL 5 to 8 h prior to collection.

### Intravitreal injections

Pigs were placed on an operating room table and pre-anesthetized with an intramuscular injection of ketamine (14 mg/kg) and acepromazine (1 mg/kg). Pigs were then anesthetized using a cone inhalation delivery system of 1-5% isoflurane. Proparacaine was administered as a topical anesthetic eyedrop to mitigate discomfort on the ocular surface. Once properly sedated, the eyes were prepared using a 5% Betadine solution (povidone-iodine solution) applied around the eye and on the ocular surface. A sterile alphonso lid speculum was placed in each eye. A sterile caliper was then used to measure 2.5 mm from the surgical limbus near 12 o’clock (modified depending on the age of the pig and the size of the eye, calculated as needed for each animal). A 31 G 5/16-inch needle on an insulin syringe was then used to inject DPBS or ASO at varying titers to have a maximum volume of 30 μL into the vitreous cavity. The needle was held in the eye for 10 seconds after injection and a sterile cotton tip applicator which had been dipped in the Betadine solution was placed over the sclera as the needle was removed to prevent reflux.

Manual intraocular pressure assessment and observation for clouding of the cornea was performed along with fundus photography using a condensing lens (20 or 28D) and an iPhone camera, Vista View, or indirect ophthalmoscope. At the completion of the procedure, antibiotic and steroid ointment was applied to the eyes.

Rats were anesthetized using a chamber inhalation delivery system of 1-5% isoflurane. Once properly sedated, animals were placed in a stereotaxic instrument. A sterile Hamilton syringe with a 33-gauge needle was placed in the stereotaxic manipulator and used to inject PBS or 5 μg of ASO bilaterally at a volume of 2 μL into the vitreous cavity.

### Electroretinogram

Pigs were tested for retinal function using a flash ERG RETeval device (LKC Technologies) as previously described (*31, 70*). Baseline ERGs were recorded either immediately before injection, or within the first week following injections to limit the time under anesthesia during injections, and subsequently every 3 months until terminal tissue harvest.

Light-adapted testing was performed after a minimum 10 min exposure to artificial light, i.e. standard operating room illumination. The rabbit/minipig, photopic 2-step light-adapted protocol was used for each eye and produced an 8.0 cd•s/m^2^ flash at 2.0 Hz followed by an 8.0 cd•s/m^2^ flicker at 28.3 Hz with a 30cd/m^2^ background (*31, 70*).

After light-adapted testing of both eyes, all lights in the room were extinguished, external light was eliminated, and a piece of blackout fabric was placed over the eyes of the pig. The RETeval device was calibrated for dark adaptation, and animals were allowed to adapt to the dark for 20 minutes. A dim red light was used as needed to set up the test.

Following dark adaptation, the rabbit/minipig, scotopic 2-step protocol was used for each eye. The first step produced an 8.0 cd•s/m^2^ flash at 0.1 Hz (dark-adapted mixed-rod and cone response), followed by a 25 cd•s/m^2^ flash at 0.05 Hz (dark-adapted mixed-rod and cone response to higher intensity flash).

Raw (unsmoothed) data values were used to calculate amplitudes. The a-wave amplitude was recorded as the pre-stimulus baseline to an a-wave trough, and the b-wave amplitude was measured from an a-wave trough to the highest waveform peak. ERG recordings were assessed by a blinded researcher and some recordings were excluded from analysis on the basis of poor electrode placement or too much noise in the waveform baseline prior to stimulation. In order to limit inter-animal variability of retinal function, ERG measurements from the eyes from individual pigs, one treated with vehicle and one with ASO, were analyzed in pair-wise comparisons and the change over time in the ERG was calculated, with the first measurement, 1 week after treatment served as baseline to account for any injection-related effects. To enable this analysis and control for pigs with vastly different ERG reading as baseline, we carried out an outlier analysis and two animals were also excluded from some analyses because ERG recordings at the start of the study were >2 SD from the group mean.

Rats were tested for retinal function using a flash ERG Celeris device (Diagnosys LLC) 4 weeks post-injection. Dark-adapted testing was performed after housing the animals in the dark room for ∼24 hours prior to testing. A 3-step dark-adapted protocol was used for each eye and produced a 0.01 cd•s/m^2^ flash at 1.0 Hz, followed by a 3.0 cd•s/m^2^ flash at 1.0 Hz, followed by a 10.0 cd•s/m^2^ flash at 1.0 Hz. Three sweeps were recorded at each intensity level per eye and the average response was used to determine the a– and b-wave amplitudes. The a-wave amplitude was recorded as the pre-stimulus baseline to an a-wave trough, and the b-wave amplitude was measured from an a-wave trough to the highest waveform peak.

### Tissue collection and processing

Following euthanasia via barbiturates (10cc/45 kg IV pentobarbital), as approved by the AVMA in SOP 2037 Swine Euthanasia, pigs were enucleated. Using a scalpel, a small slit was cut in the posterior portion of the eye near the optic nerve to isolate a piece of retinal tissue for splicing analysis. Retinal tissue was flash frozen in liquid nitrogen. Eyes were then injected with 2-3 mL of 10% formalin intravitreally through the slit and immersed in 10% formalin (20x volume of the eye) for 24 hours at room temperature. After 24 hours, the superior portion of the eye just above the lens was removed. The remaining portion of the eye was placed back into 10% formalin and fixed for another 24 hours. Following the 48 total formalin fixation time, a second cut was made to remove the inferior portion of the eye just below the optic nerve. Eyes were then placed in 50% ethanol for 1 hour, then transferred to a pathology cassette used for paraffin embedding in 70% ethanol until embedding or up to 5 days.

### Paraffin embedding

Briefly, wax pots were heated to 66°C to melt paraffin. Once melted, tissues were placed top side down in a metal wax mold and melted paraffin was poured over the tissue. Cassettes were then placed back on top of tissue and the wax was cooled on a cold plate until hardened. Tissue blocks were carefully removed and stored at room temperature until sectioned.

### Tissue Sectioning/Immunohistochemistry

Tissue blocks were sectioned at 10 µm thickness on a rotary microtome. Paraffin-sectioned tissues were processed using the Leica BOND RX Fully Automated Research Stainer (Leica Biosystems, 21.2821). Deparaffinization was performed using the Leica BOND RX Bake and Dewax protocol using Bond Dewax (Leica Biosystems, AR9222, Deer Park, IL USA). Antigen – retrieval was performed using the HIER with BOND Epitope Retrieval 2 solution (Leica Biosystems, AR9640, Deer Park, IL USA) for 20 minutes. Slides were immunolabeled using the BOND IHC IF protocol with 20-minute primary and secondary antibody incubations using 2′-O-methoxyethyl ASOs (Ionis Pharmaceuticals, 13545; 1:500) and goat anti-rabbit IgG (H+L) Alexa Fluor 647 conjugate (ThermoFisher Scientific, A-21245; 1:1000) and counterstained with Hoechst (ThermoFisher Scientific, H3570; 1:1000). Slides were cover-slipped using anti-fade fluorescence mounting medium (Abcam, Ab104135). Sections were imaged using a Nikon 90i microscope with NIS-Elements Advanced Research software (version [v.]4.20+).

### Statistical Analyses

Statistical analyses of data were performed using Prism 10.2.1 (Graphpad). Experimental sample sizes are provided in **Table S8**. Prior to analysis, ERG data were assessed by a researcher masked to treatment eye for electrode placement and eye direction on the in-unit camera system, and trace quality; poor-quality acquisitions and those with baseline measurements more than two standard deviations from the group mean were not included in statistical analyses of change from baseline due to outlier status. In assessing ERG change from baseline, one-way ANOVA with Dunnett’s multiple comparisons test was used to compare the ASO-treated and vehicle-treated *CLN3^Δ78^*eyes to vehicle-treated heterozygous controls. Contralateral ASO-treated and vehicle-treated *CLN3^Δ78^* eyes were compared using one-tailed, paired t-tests. \**P* <0.05, \*\**P* <0.01, \*\*\**P* <0.0001, \*\*\*\**P* <0.0001.

## List of Supplementary Materials

Figs S1 to S11

Tables S1 to S8

## Acknowledgements

The authors thank the staff of Exemplar Genetics, including Brian Bishop, DVM, for miniswine management and care as well as their assistance with injections, ERG, and tissue collection; Steven A. Miller for consultation on statistical methodology; Katrina A. Stratton for her insight into vision loss, its clinical manifestations, and impact on patient well-being; Estela Viera for technical support; and the members of the Hastings lab for stimulating discussions. This work is dedicated to the memory of Kiba D. Stratton, without whose lifetime of friendship, support, and love this work would not have been possible.

## Author contributions

Conceptualization: MPS, JLC, JLH, FR, PJ, JMW, MLH

Data curation: MPS, VJS, CDB, KA, MJR, FJD, HGL, JLH

Formal analysis: MPS

Funding acquisition: JMW, MLH

Investigation: MPS, VJS, WLP, KA, MJR, JLH, AVD

Methodology: MPS, JLC, VJS, WLP, PJ, FR, AVD

Project administration: MPS, TS, JMW, MLH

Resources: JLH, JMW, MLH

Software: MPS, FJD, HGL

Supervision: MPS, JMW, MLH

Validation: MPS, JLC, VJS, WLP, CDB, KA, MJR, JMW, AVD, MLH

Visualization: MPS, JLC

Writing – original draft: MPS, MLH

Writing – review & editing: MPS, JLC, VJS, WLP, CDB, KA, MJR, JLH, PJ, JMW, AVD, MLH

## Data and materials availability

The data underlying this article are available in the article and in its online supplemental material.

## Figure Legends

**Figure S1.** Molecular characterization of the *CLN3^Δ78^* minipig. (A) Sanger sequencing of gDNA from a *CLN3^Δ78^* minipig aligned with *Sus scrofa* whole genome shotgun sequence (AEMK02000016.1), illustrating the borders of the 952 bp deletion. Highlighted in green, the rAAV-mediated gene targeting generation of the *pCLN3^Δ78^* pig also resulted in a 122bp insertion from the PGK-NeoR cassette used in the process. (B) Sanger sequencing chromatogram of cDNA from a *CLN3^Δ78^* pig confirming splicing of exon 6 to exon 9. (C) PCR amplification of genomic DNA from a wild-type (+/+) or homozygote *CLN3^Δ78^* pig showing exon 7 & 8 deletion. (D) Radiolabeled RT–PCR products amplified from cDNA made from RNA isolated from retina of pig with indicated genotypes. Products are labeled on the left side of the gels. \**CLN3^Δ9^*.

**Figure S2.** ASOs induce exon 5 skipping to produce *CLN3^Δ578^* and *CLN3^Δ5678^* RNA in *CLN3^Δ78^* pig fibroblasts. (A) Radioactive RT–PCR was performed on the cDNA used in Figure 1 from RNA extracted from *CLN3^Δ78^* cells individually transfected with the indicated ASO, mock treated (C), or untreated (–). PCR was performed using primers in *CLN3* exons 4 and 10. The spliced products resulting from the amplification are labeled on the left of the gel. Below: quantification of exon 5 splicing (calculated as a percentage *CLN3^Δ5678^*/(*CLN3^Δ78^*+*CLN3^Δ5678^*)) from a single experiment shown in the gel. Cells were treated with puromycin prior to collection to block nonsense mediated decay (NMD) with the exception of cells from sample labelled “+NMD”. Size markers (bp) are shown on the right of the gel.

**Figure S3.** *CLN3* mRNA in the pig retina at different ages. (A) Radioactive RT–PCR analysis of *CLN3* in the retina of *CLN3*^+/+^ and *CLN3^Δ78^* pigs at 2, 6, 36, and 50 months (mo) of age. *HPRT1* was included as a loading control. Products are labeled on the left of the gel. Size markers (bp) are shown on the right of gel. (B) Quantification of groups shown in (A) with all labeled mRNA isoforms included in quantitation. (C) RT-PCR analysis of RNA isolated from the retina and optic nerve tissue of a heterozygote CLN3+/Δ78 pig three months after intravitreal injection ASO-16 (300 μg) into one eye and vehicle (saline, “c”) in the contralateral eye. The bands corresponding to each expected splice product are labeled. HPRT1 was analyzed as a control.

**Figure S4.** ASO-16 is well-tolerated at 80 μg in the retina of *CLN3^Δ78^* minipigs. ERG raw amplitudes of a-wave, b-wave, and 28.3 Hz flicker at (A) on the day of and (B) 1 week post injection of DPBS or with 80 μg of ASO-16. Bars show s.e.m.; one-way ANOVA with Dunnett’s multiple comparisons; \**P* <0.05, \*\**P* <0.01

**Figure S5.** No changes in retinal conduction rate observed in *CLN3^Δ78^* minipigs. The change in ERG (A-C) a-wave implicit time, (D-F) b-wave implicit time, and (G) implicit time from harmonics is visualized comparing the ASO-treated and vehicle-treated eye of *CLN3^Δ78^* pigs over time. Bars show s.e.m.; one-tailed paired t-tests; \**P* <0.05.

**Figure S6.** High responder pig ERG waveforms. (A-D) Comparison of ERG waveforms from the ASO-treated and vehicle-treated eye of four *CLN3^Δ78^*pigs identified as high responders across the (A) 28.3 Hz flicker (8.0 cd•s/m^2^), (B) light-adapted bright flash (8.0 cd•s/m^2^), (C) dark-adapted bright flash (8.0 cd•s/m^2^), (D) or dark-adapted super bright flash (25.0 cd•s/m^2^) 9 months post-treatment.

**Figure S7.** Non-responder ERG waveforms. (A-D) Comparison of ERG waveforms from the ASO-treated (blue) and vehicle-treated (red) eye of *CLN3^Δ78^* pigs who did not show improvement in retinal function across the (A) 28.3 Hz flicker (8.0 cd•s/m^2^), (B) light-adapted bright flash (8.0 cd•s/m^2^), (C) dark-adapted bright flash (8.0 cd•s/m^2^), (D) or dark-adapted super bright flash (25.0 cd•s/m^2^) 9 months post-treatment. (NR) Poor recordings were omitted due to background noise making the waveforms unreliable.

**Figure S8.** Intravitreal delivery of ASO-16 improves retinal function in a subset of high-responding pigs. (A-G) Comparison of the raw ERG amplitudes of individual *CLN3^Δ78^* pigs 9 months post-treatment (A-C) a-wave, (D-F) b-wave, and (G) amplitudes at 28.3 Hz flicker. Comparing vehicle-treated and 80 µg ASO-16 treated eyes within each *CLN3^Δ78^* pig with bright flash (8.0 cd•s/m^2^) or super bright flash (25.0 cd•s/m^2^). Connecting lines indicate which data points correspond to contralateral eyes within the same animal, with green lines, highlighting high-responding animals. One-tailed paired t-test comparisons for all animals shown in black; \**P* <0.05; one-tailed paired t-test comparisons for high-responding animals shown in green lettering; ^#^P<0.05, ^##^P<0.01.

**Figure S9.** Retinal function and splicing 12 months after a single treatment with ASO-16. (A-G) The change from baseline (CFB) in ERG amplitudes of individual *CLN3^Δ78^*pigs 12 months post-treatment (A-C) a-wave (predominately photoreceptors), (D-F) b-wave (predominately bipolar cells), and (G) amplitudes at 28.3 Hz flicker (cone bipolar cells only). Comparing vehicle-treated and 80 µg ASO-16 treated eyes within each *CLN3^Δ78^* pig with bright flash (8.0 cd•s/m^2^) or super bright flash (25.0 cd•s/m^2^). Connecting lines indicate which data points correspond to contralateral eyes within the same animal, with green lines highlighting high-responding animals from 9 months post-treatment. One-tailed paired t-test comparisons for all animals shown in black; one-tailed paired t-test comparisons for high-responding animals shown in green; ^#^P<0.05. (H) Products of radioactive RT–PCR analysis of pig retinal RNA isolated from three homozygous *CLN3^Δ78^* pigs a year after intravitreal injection of DPBS vehicle (control) into one eye and ASO-16 (80 µg) into the other eye separated by PAGE. *HPRT1* was included as a loading control. Products are labeled on the left of the gel. Size markers (bp) are shown on the right of gel. (I) Quantification of exon 5 splicing (calculated as a percentage *CLN3^Δ578^*/(*CLN3^Δ78^*+*CLN3^Δ578^*)x100. \*\**P* <0.01; paired t-test.

**Figure S10.** A single intravitreal injection of ASO-16 and ASO-29a is well tolerated in rats. Dark-adapted ERG of the a-wave amplitude (A) and b-wave amplitude (B) comparing the ASO-treated and vehicle-treated eye of wildtype rats 4 weeks post-treatment after the indicated form of stimulation. Bars show s.e.m.; one-way ANOVA comparing ASO-16 and ASO-29a to control treated.

**Figure S11.** An 80 μg dose of ASO-29a is well-tolerated in the retina of *CLN3^Δ78^* minipigs. ERG raw amplitudes of a-wave, b-wave, and 28.3 Hz flicker following bright (8.0 cd•s/m^2^) and super bright (25.0 cd•s/m^2^) light stimuli in a light adapted or dark-adapted environment at (A) 1 week, (B) 3 months, and (C) 6 months post injection of DPBS or with either 80 μg or 160 μg of ASO-29a. Bars show s.e.m.; one-way ANOVA with Dunnett’s multiple comparisons of vehicle treated to 80 μg and 160 μg ASO-29a treated; \**P* <0.05, \*\*\**P* <0.001.

## References and Notes

1. P. B. Munroe, H. M. Mitchison, A. M. O’Rawe, J. W. Anderson, R. M. Boustany, T. J. Lerner, P. E. Taschner, N. de Vos, M. H. Breuning, R. M. Gardiner, S. E. Mole, Spectrum of mutations in the Batten disease gene, CLN3. Am J Hum Genet 61, 310–316 (1997).

2. D. E. Sleat, E. Gedvilaite, Y. Zhang, P. Lobel, J. Xing, Analysis of large-scale whole exome sequencing data to determine the prevalence of genetically-distinct forms of neuronal ceroid lipofuscinosis. Gene 593, 284–291 (2016).

3. Isolation of a novel gene underlying Batten disease, CLN3. The International Batten Disease Consortium. Cell 82, 949–957 (1995).

4. A. Schulz, N. Patel, J. J. Brudvig, F. Stehr, J. M. Weimer, E. F. Augustine, The parent and family impact of CLN3 disease: an observational survey-based study. Orphanet J Rare Dis 19, 125 (2024).

5. M. C. Masten, J. D. Williams, J. Vermilion, H. R. Adams, A. Vierhile, A. Collins, F. J. Marshall, E. F. Augustine, J. W. Mink, The CLN3 Disease Staging System: A new tool for clinical research in Batten disease. Neurology 94, e2436–e2440 (2020).

6. J. W. Mink, E. F. Augustine, H. R. Adams, F. J. Marshall, J. M. Kwon, Classification and Natural History of the Neuronal Ceroid Lipofuscinoses. Journal of Child Neurology 28, 1101–1105 (2013).

7. A. K. Nielsen, J. R. Ostergaard, Do females with juvenile ceroid lipofuscinosis (Batten disease) have a more severe disease course? The Danish experience. Eur J Paediatr Neurol 17, 265–268 (2013).

8. J. Cialone, H. Adams, E. F. Augustine, F. J. Marshall, J. M. Kwon, N. Newhouse, A. Vierhile, E. Levy, L. S. Dure, K. R. Rose, D. Ramirez-Montealegre, E. A. de Blieck, J. W. Mink, Females experience a more severe disease course in Batten disease. J Inherit Metab Dis 35, 549–555 (2012).

9. J. Collins, G. E. Holder, H. Herbert, G. G. Adams, Batten disease: features to facilitate early diagnosis. Br J Ophthalmol 90, 1119–1124 (2006).

10. G. A. Wright, M. Georgiou, A. G. Robson, N. Ali, A. Kalhoro, S. K. Holthaus, N. Pontikos, N. Oluonye, E. R. de Carvalho, M. M. Neveu, R. G. Weleber, M. Michaelides, Juvenile Batten Disease (CLN3): Detailed Ocular Phenotype, Novel Observations, Delayed Diagnosis, Masquerades, and Prospects for Therapy. Ophthalmol Retina 4, 433–445 (2020).

11. R. G. Weleber, The dystrophic retina in multisystem disorders: the electroretinogram in neuronal ceroid lipofuscinoses. Eye (Lond) 12 **(Pt** **3b**), 580–590 (1998).

12. D. G. Birch, Retinal degeneration in retinitis pigmentosa and neuronal ceroid lipofuscinosis: An overview. Mol Genet Metab 66, 356–366 (1999).

13. L. B. Eksandh, V. B. Ponjavic, P. B. Munroe, H. E. Eiberg, P. E. Uvebrant, B. E. Ehinger, S. E. Mole, S. Andreasson, Full-field ERG in patients with Batten/Spielmeyer-Vogt disease caused by mutations in the CLN3 gene. Ophthalmic Genet 21, 69–77 (2000).

14. C. J. Minnis, S. Townsend, J. Petschnigg, E. Tinelli, J. Bahler, C. Russell, S. E. Mole, Global network analysis in Schizosaccharomyces pombe reveals three distinct consequences of the common 1-kb deletion causing juvenile CLN3 disease. Sci Rep 11, 6332 (2021).

15. A. Petcherski, U. Chandrachud, E. S. Butz, M. C. Klein, W. N. Zhao, S. A. Reis, S. J. Haggarty, M. O. Ruonala, S. L. Cotman, An Autophagy Modifier Screen Identifies Small Molecules Capable of Reducing Autophagosome Accumulation in a Model of CLN3-Mediated Neurodegeneration. Cells 8, (2019).

16. J. L. Centa, F. M. Jodelka, A. J. Hinrich, T. B. Johnson, J. Ochaba, M. Jackson, D. M. Duelli, J. M. Weimer, F. Rigo, M. L. Hastings, Therapeutic efficacy of antisense oligonucleotides in mouse models of CLN3 Batten disease. Nat Med 26, 1444–1451 (2020).

17. A. Kyttala, G. Ihrke, J. Vesa, M. J. Schell, J. P. Luzio, Two motifs target Batten disease protein CLN3 to lysosomes in transfected nonneuronal and neuronal cells. Mol Biol Cell 15, 1313–1323 (2004).

18. J. N. Miller, C. H. Chan, D. A. Pearce, The role of nonsense-mediated decay in neuronal ceroid lipofuscinosis. Hum Mol Genet 22, 2723–2734 (2013).

19. S. E. Mole, The genetic spectrum of human neuronal ceroid-lipofuscinoses. Brain Pathol 14, 70–76 (2004).

20. E. Kida, W. Kaczmarski, A. A. Golabek, A. Kaczmarski, M. Michalewski, K. E. Wisniewski, Analysis of intracellular distribution and trafficking of the CLN3 protein in fusion with the green fluorescent protein in vitro. Mol Genet Metab 66, 265–271 (1999).

21. K. F. Tung, C. Y. Pan, C. H. Chen, W. C. Lin, Top-ranked expressed gene transcripts of human protein-coding genes investigated with GTEx dataset. Sci Rep 10, 16245 (2020).

22. H. Y. Zhang, C. Minnis, E. Gustavsson, M. Ryten, S. E. Mole, CLN3 transcript complexity revealed by long-read RNA sequencing analysis. BMC Med Genomics 17, 244 (2024).

23. F. Cunningham, J. E. Allen, J. Allen, J. Alvarez-Jarreta, M. R. Amode, I. M. Armean, O. Austine-Orimoloye, A. G. Azov, I. Barnes, R. Bennett, A. Berry, J. Bhai, A. Bignell, K. Billis, S. Boddu, L. Brooks, M. Charkhchi, C. Cummins, L. Da Rin Fioretto, C. Davidson, K. Dodiya, S. Donaldson, B. El Houdaigui, T. El Naboulsi, R. Fatima, C. G. Giron, T. Genez, J. G. Martinez, C. Guijarro-Clarke, A. Gymer, M. Hardy, Z. Hollis, T. Hourlier, T. Hunt, T. Juettemann, V. Kaikala, M. Kay, I. Lavidas, T. Le, D. Lemos, J. C. Marugan, S. Mohanan, A. Mushtaq, M. Naven, D. N. Ogeh, A. Parker, A. Parton, M. Perry, I. Pilizota, I. Prosovetskaia, M. P. Sakthivel, A. I. A. Salam, B. M. Schmitt, H. Schuilenburg, D. Sheppard, J. G. Perez-Silva, W. Stark, E. Steed, K. Sutinen, R. Sukumaran, D. Sumathipala, M. M. Suner, M. Szpak, A. Thormann, F. F. Tricomi, D. Urbina-Gomez, A. Veidenberg, T. A. Walsh, B. Walts, N. Willhoft, A. Winterbottom, E. Wass, M. Chakiachvili, B. Flint, A. Frankish, S. Giorgetti, L. Haggerty, S. E. Hunt, I. I. Gr, J. E. Loveland, F. J. Martin, B. Moore, J. M. Mudge, M. Muffato, E. Perry, M. Ruffier, J. Tate, D. Thybert, S. J. Trevanion, S. Dyer, P. W. Harrison, K. L. Howe, A. D. Yates, D. R. Zerbino, P. Flicek, Ensembl 2022. Nucleic Acids Res 50, D988–D995 (2022).

24. N. A. O’Leary, M. W. Wright, J. R. Brister, S. Ciufo, D. Haddad, R. McVeigh, B. Rajput, B. Robbertse, B. Smith-White, D. Ako-Adjei, A. Astashyn, A. Badretdin, Y. Bao, O. Blinkova, V. Brover, V. Chetvernin, J. Choi, E. Cox, O. Ermolaeva, C. M. Farrell, T. Goldfarb, T. Gupta, D. Haft, E. Hatcher, W. Hlavina, V. S. Joardar, V. K. Kodali, W. Li, D. Maglott, P. Masterson, K. M. McGarvey, M. R. Murphy, K. O’Neill, S. Pujar, S. H. Rangwala, D. Rausch, L. D. Riddick, C. Schoch, A. Shkeda, S. S. Storz, H. Sun, F. Thibaud-Nissen, I. Tolstoy, R. E. Tully, A. R. Vatsan, C. Wallin, D. Webb, W. Wu, M. J. Landrum, A. Kimchi, T. Tatusova, M. DiCuccio, P. Kitts, T. D. Murphy, K. D. Pruitt, Reference sequence (RefSeq) database at NCBI: current status, taxonomic expansion, and functional annotation. Nucleic Acids Res 44, D733–745 (2016).

25. M. A. Havens, M. L. Hastings, Splice-switching antisense oligonucleotides as therapeutic drugs. Nucleic Acids Res 44, 6549–6563 (2016).

26. K. Dulla, M. Aguila, A. Lane, K. Jovanovic, D. A. Parfitt, I. Schulkens, H. L. Chan, I. Schmidt, W. Beumer, L. Vorthoren, R. W. J. Collin, A. Garanto, L. Duijkers, A. Brugulat-Panes, M. Semo, A. A. Vugler, P. Biasutto, P. Adamson, M. E. Cheetham, Splice-Modulating Oligonucleotide QR-110 Restores CEP290 mRNA and Function in Human c.2991+1655A>G LCA10 Models. Mol Ther Nucleic Acids 12, 730–740 (2018).

27. S. R. Russell, A. V. Drack, A. V. Cideciyan, S. G. Jacobson, B. P. Leroy, C. Van Cauwenbergh, A. C. Ho, A. V. Dumitrescu, I. C. Han, M. Martin, W. L. Pfeifer, E. H. Sohn, J. Walshire, A. V. Garafalo, A. K. Krishnan, C. A. Powers, A. Sumaroka, A. J. Roman, E. Vanhonsebrouck, E. Jones, F. Nerinckx, J. De Zaeytijd, R. W. J. Collin, C. Hoyng, P. Adamson, M. E. Cheetham, M. R. Schwartz, W. den Hollander, F. Asmus, G. Platenburg, D. Rodman, A. Girach, Intravitreal antisense oligonucleotide sepofarsen in Leber congenital amaurosis type 10: a phase 1b/2 trial. Nat Med 28, 1014–1021 (2022).

28. K. Dulla, R. Slijkerman, H. C. van Diepen, S. Albert, M. Dona, W. Beumer, J. J. Turunen, H. L. Chan, I. A. Schulkens, L. Vorthoren, C. den Besten, L. Buil, I. Schmidt, J. Miao, H. Venselaar, J. Zang, S. C. F. Neuhauss, T. Peters, S. Broekman, R. Pennings, H. Kremer, G. Platenburg, P. Adamson, E. de Vrieze, E. van Wijk, Antisense oligonucleotide-based treatment of retinitis pigmentosa caused by USH2A exon 13 mutations. Mol Ther 29, 2441–2455 (2021).

29. J. Grainok, I. L. Pitout, F. K. Chen, S. McLenachan, R. C. Heath Jeffery, C. Mitrpant, S. Fletcher, A Precision Therapy Approach for Retinitis Pigmentosa 11 Using Splice-Switching Antisense Oligonucleotides to Restore the Open Reading Frame of PRPF31. Int J Mol Sci 25, (2024).

30. J. L. Centa, M. P. Stratton, M. A. Pratt, J. R. Osterlund Oltmanns, D. G. Wallace, S. A. Miller, J. M. Weimer, M. L. Hastings, Protracted CLN3 Batten disease in mice that genetically model an exon-skipping therapeutic approach. Mol Ther Nucleic Acids 33, 15–27 (2023).

31. V. J. Swier, K. A. White, T. B. Johnson, X. Wang, J. Han, D. A. Pearce, R. Singh, A. V. Drack, W. Pfeifer, C. S. Rogers, J. J. Brudvig, J. M. Weimer, A novel porcine model of CLN3 Batten disease recapitulates clinical phenotypes. Dis Model Mech 16, (2023).

32. C. Kostic, Y. Arsenijevic, Animal modelling for inherited central vision loss. J Pathol 238, 300–310 (2016).

33. J. M. Weimer, A. W. Custer, J. W. Benedict, N. A. Alexander, E. Kingsley, H. J. Federoff, J. D. Cooper, D. A. Pearce, Visual deficits in a mouse model of Batten disease are the result of optic nerve degeneration and loss of dorsal lateral geniculate thalamic neurons. Neurobiol Dis 22, 284–293 (2006).

34. G. M. Seigel, A. Lotery, A. Kummer, D. J. Bernard, N. D. Greene, M. Turmaine, T. Derksen, R. L. Nussbaum, B. Davidson, J. Wagner, H. M. Mitchison, Retinal pathology and function in a Cln3 knockout mouse model of juvenile Neuronal Ceroid Lipofuscinosis (batten disease). Mol Cell Neurosci 19, 515–527 (2002).

35. H. N. Schwahn, F. Gekeler, K. Kohler, K. Kobuch, H. G. Sachs, F. Schulmeyer, W. Jakob, V. P. Gabel, E. Zrenner, Studies on the feasibility of a subretinal visual prosthesis: data from Yucatan micropig and rabbit. Graefes Arch Clin Exp Ophthalmol 239, 961–967 (2001).

36. F. Ghosh, K. Engelsberg, R. V. English, R. M. Petters, Long-term neuroretinal full-thickness transplants in a large animal model of severe retinitis pigmentosa. Graefes Arch Clin Exp Ophthalmol 245, 835–846 (2007).

37. H. Klassen, J. F. Kiilgaard, T. Zahir, B. Ziaeian, I. Kirov, E. Scherfig, K. Warfvinge, M. J. Young, Progenitor cells from the porcine neural retina express photoreceptor markers after transplantation to the subretinal space of allorecipients. Stem Cells 25, 1222–1230 (2007).

38. C. Mussolino, M. della Corte, S. Rossi, F. Viola, U. Di Vicino, E. Marrocco, S. Neglia, M. Doria, F. Testa, R. Giovannoni, M. Crasta, M. Giunti, E. Villani, M. Lavitrano, M. L. Bacci, R. Ratiglia, F. Simonelli, A. Auricchio, E. M. Surace, AAV-mediated photoreceptor transduction of the pig cone-enriched retina. Gene Ther 18, 637–645 (2011).

39. S. M. Shrader, W. F. Greentree, Gottingen Minipigs in Ocular Research. Toxicol Pathol 46, 403–407 (2018).

40. S. P. Henry, R. C. Miner, W. L. Drew, J. Fitchett, C. York-Defalco, L. M. Rapp, A. A. Levin, Antiviral activity and ocular kinetics of antisense oligonucleotides designed to inhibit CMV replication. Invest Ophthalmol Vis Sci 42, 2646–2651 (2001).

41. M. Byrne, V. Vathipadiekal, L. Apponi, N. Iwamoto, P. Kandasamy, K. Longo, F. Liu, R. Looby, L. Norwood, A. Shah, J. D. Shelke, C. Shivalila, H. Yang, Y. Yin, L. Guo, K. Bowman, C. Vargeese, Stereochemistry Enhances Potency, Efficacy, and Durability of Malat1 Antisense Oligonucleotides In Vitro and In Vivo in Multiple Species. Transl Vis Sci Technol 10, 23 (2021).

42. A. Shafiee, G. L. McIntire, L. C. Sidebotham, K. W. Ward, Experimental determination and allometric prediction of vitreous volume, and retina and lens weights in Gottingen minipigs. Vet Ophthalmol 11, 193–196 (2008).

43. in Webvision: The Organization of the Retina and Visual System, H. Kolb, E. Fernandez, R. Nelson, Eds. (Salt Lake City (UT), 1995).

44. A. V. Cideciyan, S. G. Jacobson, A. V. Drack, A. C. Ho, J. Charng, A. V. Garafalo, A. J. Roman, A. Sumaroka, I. C. Han, M. D. Hochstedler, W. L. Pfeifer, E. H. Sohn, M. Taiel, M. R. Schwartz, P. Biasutto, W. Wit, M. E. Cheetham, P. Adamson, D. M. Rodman, G. Platenburg, M. D. Tome, I. Balikova, F. Nerinckx, J. Zaeytijd, C. Van Cauwenbergh, B. P. Leroy, S. R. Russell, Effect of an intravitreal antisense oligonucleotide on vision in Leber congenital amaurosis due to a photoreceptor cilium defect. Nat Med 25, 225–228 (2019).

45. M. M. Ouseph, M. E. Kleinman, Q. J. Wang, Vision loss in juvenile neuronal ceroid lipofuscinosis (CLN3 disease). Ann N Y Acad Sci 1371, 55–67 (2016).

46. H. H. Goebel, J. D. Fix, W. Zeman, The fine structure of the retina in neuronal ceroid-lipofuscinosis. Am J Ophthalmol 77, 25–39 (1974).

47. C. Tang, J. Han, S. Dalvi, K. Manian, L. Winschel, S. Volland, C. A. Soto, C. A. Galloway, W. Spencer, M. Roll, C. Milliner, V. L. Bonilha, T. B. Johnson, L. Latchney, J. M. Weimer, E. F. Augustine, J. W. Mink, V. K. Gullapalli, M. Chung, D. S. Williams, R. Singh, A human model of Batten disease shows role of CLN3 in phagocytosis at the photoreceptor-RPE interface. Commun Biol 4, 161 (2021).

48. S. M. Kleine Holthaus, M. Aristorena, R. Maswood, O. Semenyuk, J. Hoke, A. Hare, A. J. Smith, S. E. Mole, R. R. Ali, Gene Therapy Targeting the Inner Retina Rescues the Retinal Phenotype in a Mouse Model of CLN3 Batten Disease. Hum Gene Ther 31, 709–718 (2020).

49. A. Girach, I. Audo, D. G. Birch, R. M. Huckfeldt, B. L. Lam, B. P. Leroy, M. Michaelides, S. R. Russell, J. M. F. Sallum, K. Stingl, S. H. Tsang, P. Yang, RNA-based therapies in inherited retinal diseases. Ther Adv Ophthalmol 14, 25158414221134602 (2022).

50. D. Dalkara, O. Goureau, K. Marazova, J. A. Sahel, Let There Be Light: Gene and Cell Therapy for Blindness. Hum Gene Ther 27, 134–147 (2016).

51. A. Gupta, K. N. Kafetzis, A. D. Tagalakis, C. Yu-Wai-Man, RNA therapeutics in ophthalmology – translation to clinical trials. Exp Eye Res 205, 108482 (2021).

52. A. Grzybowski, R. Told, S. Sacu, F. Bandello, E. Moisseiev, A. Loewenstein, U. Schmidt-Erfurth, B. Euretina, 2018 Update on Intravitreal Injections: Euretina Expert Consensus Recommendations. Ophthalmologica 239, 181–193 (2018).

53. X. Gerard, I. Perrault, A. Munnich, J. Kaplan, J. M. Rozet, Intravitreal Injection of Splice-switching Oligonucleotides to Manipulate Splicing in Retinal Cells. Mol Ther Nucleic Acids 4, e250 (2015).

54. A. A. Aref, Management of immediate and sustained intraocular pressure rise associated with intravitreal antivascular endothelial growth factor injection therapy. Curr Opin Ophthalmol 23, 105–110 (2012).

55. A. V. Cideciyan, S. G. Jacobson, A. C. Ho, M. Swider, A. Sumaroka, A. J. Roman, V. Wu, R. C. Russell, I. Viarbitskaya, A. V. Garafalo, M. R. Schwartz, A. Girach, Durable vision improvement after a single intravitreal treatment with antisense oligonucleotide in CEP290-LCA: Replication in two eyes. Am J Ophthalmol Case Rep 32, 101873 (2023).

56. T. Dangouloff, L. Servais, Clinical Evidence Supporting Early Treatment Of Patients With Spinal Muscular Atrophy: Current Perspectives. Ther Clin Risk Manag 15, 1153–1161 (2019).

57. D. C. De Vivo, E. Bertini, K. J. Swoboda, W. L. Hwu, T. O. Crawford, R. S. Finkel, J. Kirschner, N. L. Kuntz, J. A. Parsons, M. M. Ryan, R. J. Butterfield, H. Topaloglu, T. Ben-Omran, V. A. Sansone, Y. J. Jong, F. Shu, J. F. Staropoli, D. Kerr, A. W. Sandrock, C. Stebbins, M. Petrillo, G. Braley, K. Johnson, R. Foster, S. Gheuens, I. Bhan, S. P. Reyna, S. Fradette, W. Farwell, N. S. Group, Nusinersen initiated in infants during the presymptomatic stage of spinal muscular atrophy: Interim efficacy and safety results from the Phase 2 NURTURE study. Neuromuscul Disord 29, 842–856 (2019).

58. M. A. Pane, K. L. Puranam, R. M. Boustany, Expression of cln3 in human NT2 neuronal precursor cells and neonatal rat brain. Pediatr Res 46, 367–374 (1999).

59. S. F. Murray, A. Jazayeri, M. T. Matthes, D. Yasumura, H. Yang, R. Peralta, A. Watt, S. Freier, G. Hung, P. S. Adamson, S. Guo, B. P. Monia, M. M. LaVail, M. L. McCaleb, Allele-Specific Inhibition of Rhodopsin With an Antisense Oligonucleotide Slows Photoreceptor Cell Degeneration. Invest Ophthalmol Vis Sci 56, 6362–6375 (2015).

60. M. J. Chandler, P. J. Smith, D. A. Samuelson, E. O. MacKay, Photoreceptor density of the domestic pig retina. Vet Ophthalmol 2, 179–184 (1999).

61. A. Hendrickson, K. Bumsted-O’Brien, R. Natoli, V. Ramamurthy, D. Possin, J. Provis, Rod photoreceptor differentiation in fetal and infant human retina. Exp Eye Res 87, 415–426 (2008).

62. V. Vrolyk, M. J. Desmarais, D. Lambert, J. Haruna, M. O. Benoit-Biancamano, Neonatal and Juvenile Ocular Development in Gottingen Minipigs and Domestic Pigs: A Histomorphological and Immunohistochemical Study. Vet Pathol 57, 889–914 (2020).

63. J. Guduric-Fuchs, L. J. Ringland, P. Gu, M. Dellett, D. B. Archer, T. Cogliati, Immunohistochemical study of pig retinal development. Mol Vis 15, 1915–1928 (2009).

64. S. J. Murray, K. N. Russell, T. R. Melzer, S. J. Gray, S. J. Heap, D. N. Palmer, N. L. Mitchell, Intravitreal gene therapy protects against retinal dysfunction and degeneration in sheep with CLN5 Batten disease. Exp Eye Res 207, 108600 (2021).

65. S. J. Murray, M. P. Wellby, G. K. Barrell, K. N. Russell, A. R. Deane, J. R. Wynyard, S. J. Gray, D. N. Palmer, N. L. Mitchell, Efficacy of dual intracerebroventricular and intravitreal CLN5 gene therapy in sheep prompts the first clinical trial to treat CLN5 Batten disease. Front Pharmacol 14, 1212235 (2023).

66. F. Corpet, Multiple sequence alignment with hierarchical clustering. Nucleic Acids Res 16, 10881–10890 (1988).

67. B. F. Baker, S. S. Lot, T. P. Condon, S. Cheng-Flournoy, E. A. Lesnik, H. M. Sasmor, C. F. Bennett, 2’-O-(2-Methoxy)ethyl-modified anti-intercellular adhesion molecule 1 (ICAM-1) oligonucleotides selectively increase the ICAM-1 mRNA level and inhibit formation of the ICAM-1 translation initiation complex in human umbilical vein endothelial cells. J Biol Chem 272, 11994–12000 (1997).

68. F. Rigo, S. J. Chun, D. A. Norris, G. Hung, S. Lee, J. Matson, R. A. Fey, H. Gaus, Y. Hua, J. S. Grundy, A. R. Krainer, S. P. Henry, C. F. Bennett, Pharmacology of a central nervous system delivered 2’-O-methoxyethyl-modified survival of motor neuron splicing oligonucleotide in mice and nonhuman primates. J Pharmacol Exp Ther 350, 46–55 (2014).

69. E. E. Swayze, A. M. Siwkowski, E. V. Wancewicz, M. T. Migawa, T. K. Wyrzykiewicz, G. Hung, B. P. Monia, C. F. Bennett, Antisense oligonucleotides containing locked nucleic acid improve potency but cause significant hepatotoxicity in animals. Nucleic Acids Res 35, 687–700 (2007).

70. V. J. Swier, K. A. White, T. B. Johnson, J. C. Sieren, H. J. Johnson, K. Knoernschild, X. Wang, F. A. Rohret, C. S. Rogers, D. A. Pearce, J. J. Brudvig, J. M. Weimer, A Novel Porcine Model of CLN2 Batten Disease that Recapitulates Patient Phenotypes. Neurotherapeutics 19, 1905–1919 (2022).

