## Supplemental Figures & Tables for "Therapeutic antisense oligonucleotide mitigates retinal dysfunction in a pig model of CLN3 Batten disease"

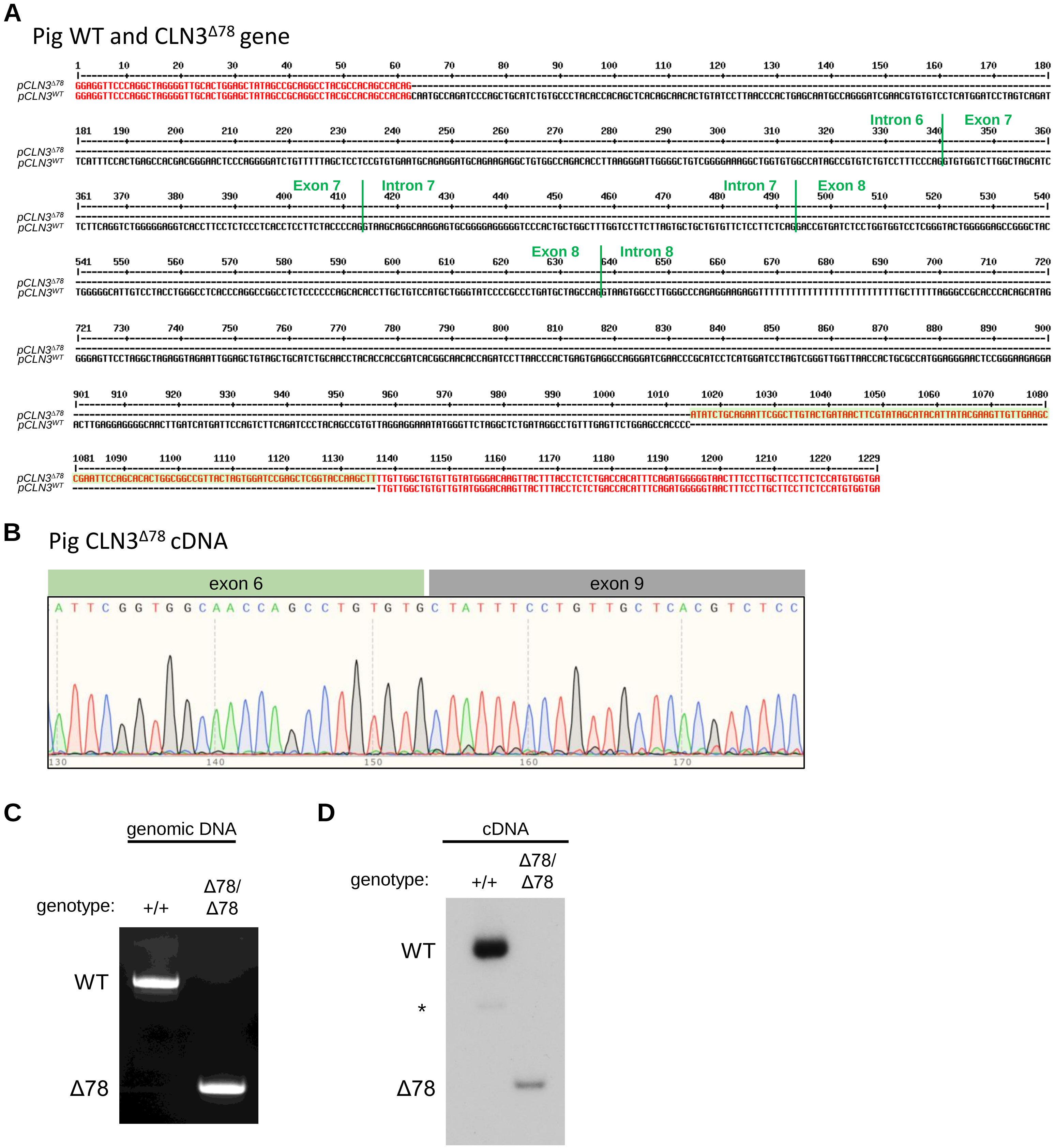
A
Pig WT and CLN3Δ78 gene
pCLN3Δ78
pCLN3WT
Intron 6
Exon 7
Intron 7
Exon 8
Intron 8
B
Pig CLN3Δ78 cDNA
exon 6
exon 9
C
genomic DNA
genotype:
+/+
Δ78/
Δ78
WT
Δ78
D
cDNA
genotype:
+/+
Δ78/
Δ78
WT
*
Δ78

**Figure S1**

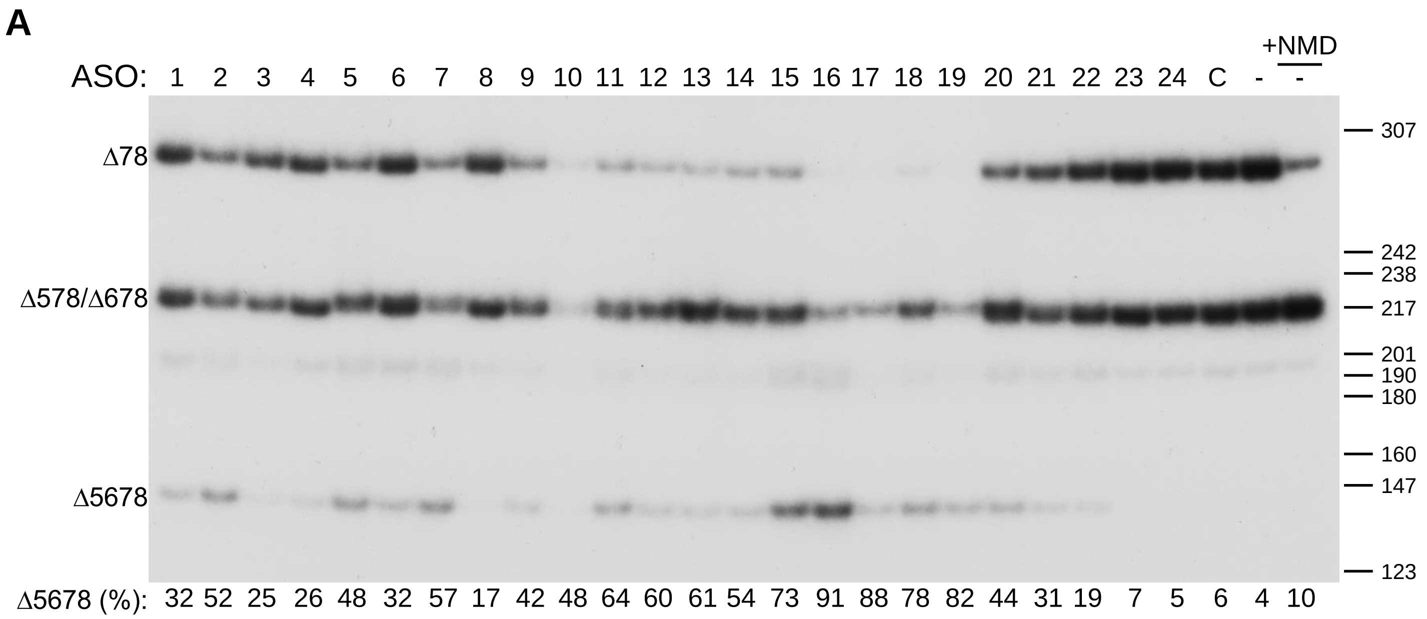
A
+NMD
ASO: 1 2 3 4 5 6 7 8 9 10 11 12 13 14 15 16 17 18 19 20 21 22 23 24 C - -
Δ78
Δ578/Δ678
Δ5678
307
242
238
217
201
190
180
160
147
123
Δ5678 (%): 32 52 25 26 48 32 57 17 42 48 64 60 61 54 73 91 88 78 82 44 31 19 7 5 6 4 10

**Figure S2**

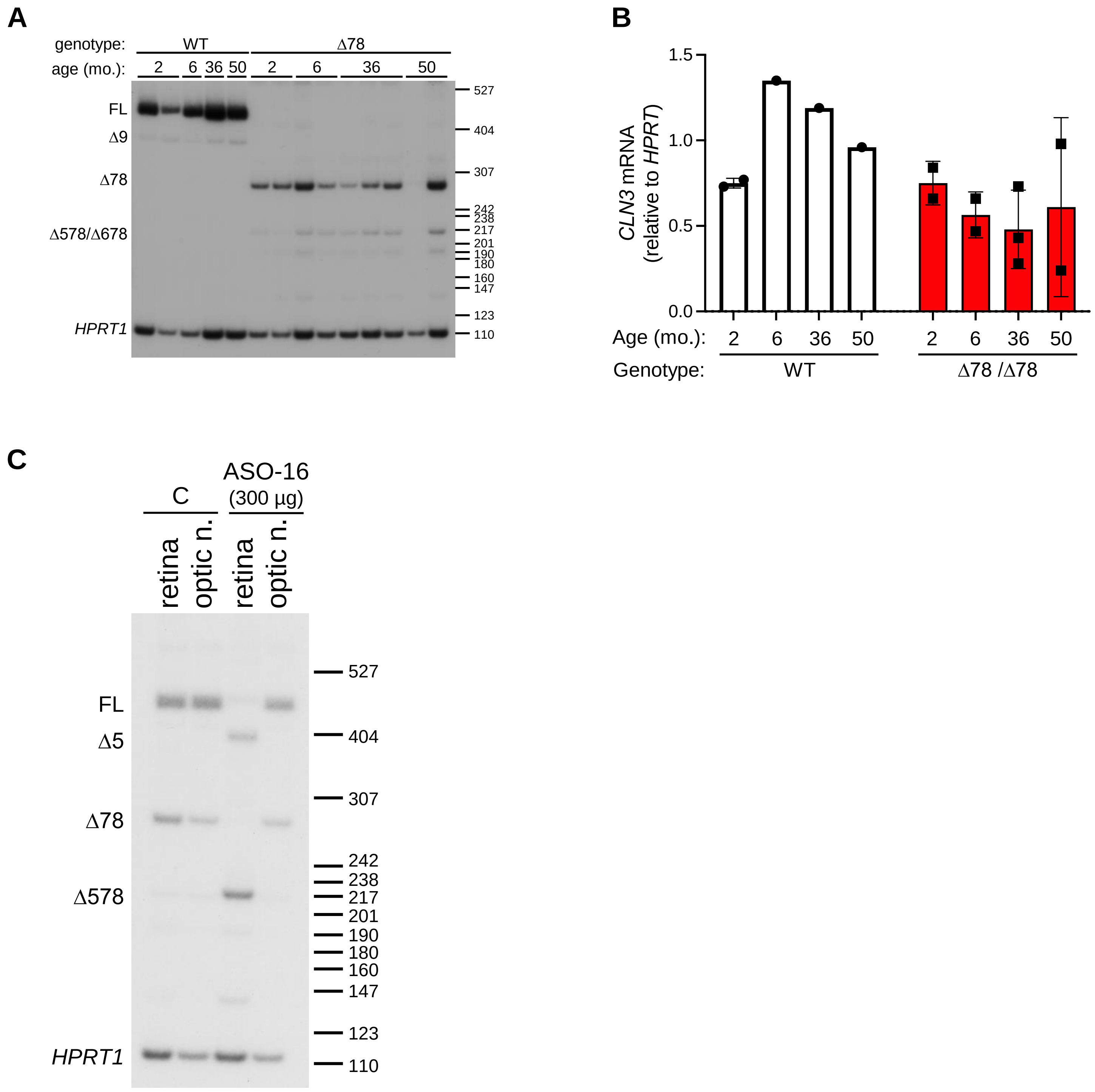

A
genotype:
WT
Δ78
age (mo.):
2
6
36
50
2
6
36
50
FL
Δ9
Δ78
Δ578/Δ678
HPRT1
527
404
307
242
238
217
201
190
180
160
147
123
110
B
CLN3 mRNA (relative to HPRT)
1.5
1.0
0.5
0.0
Age (mo.):
2
6
36
50
2
6
36
50
Genotype:
WT
Δ78 /Δ78
C
C
ASO-16 (300 μg)
retina
optic n.
retina
optic n.
FL
Δ5
Δ78
Δ578
HPRT1
527
404
307
242
238
217
201
190
180
160
147
123
110


**Figure S3**

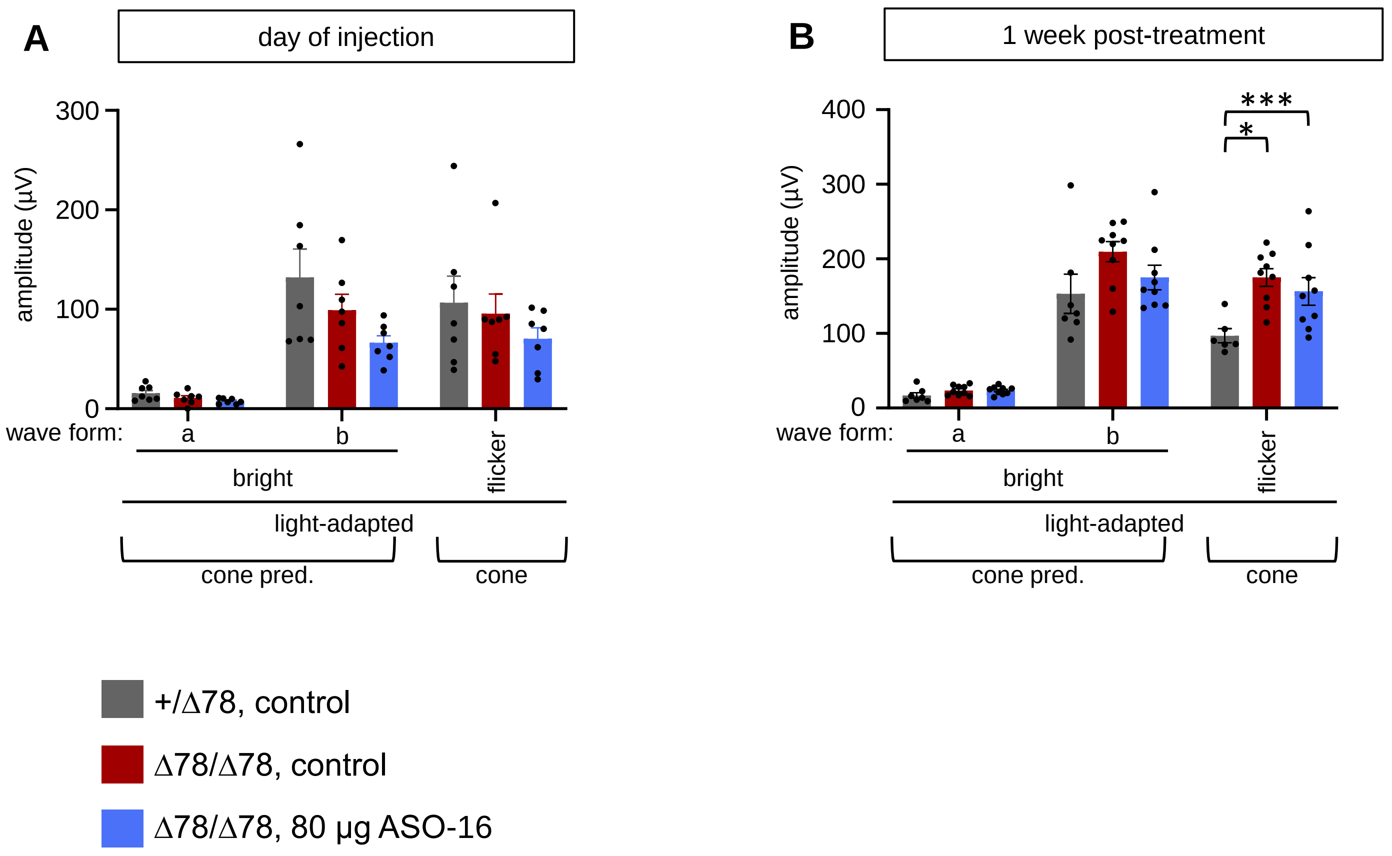
A
day of injection
300
200
100
0
amplitude (µV)
wave form:
a
b
flicker
bright
light-adapted
cone pred.
cone
B
1 week post-treatment
400
300
200
100
0
amplitude (µV)
***
*
wave form:
a
b
flicker
bright
light-adapted
cone pred.
cone
+/Δ78, control
Δ78/Δ78, control
Δ78/Δ78, 80 µg ASO-16

**Figure S4**

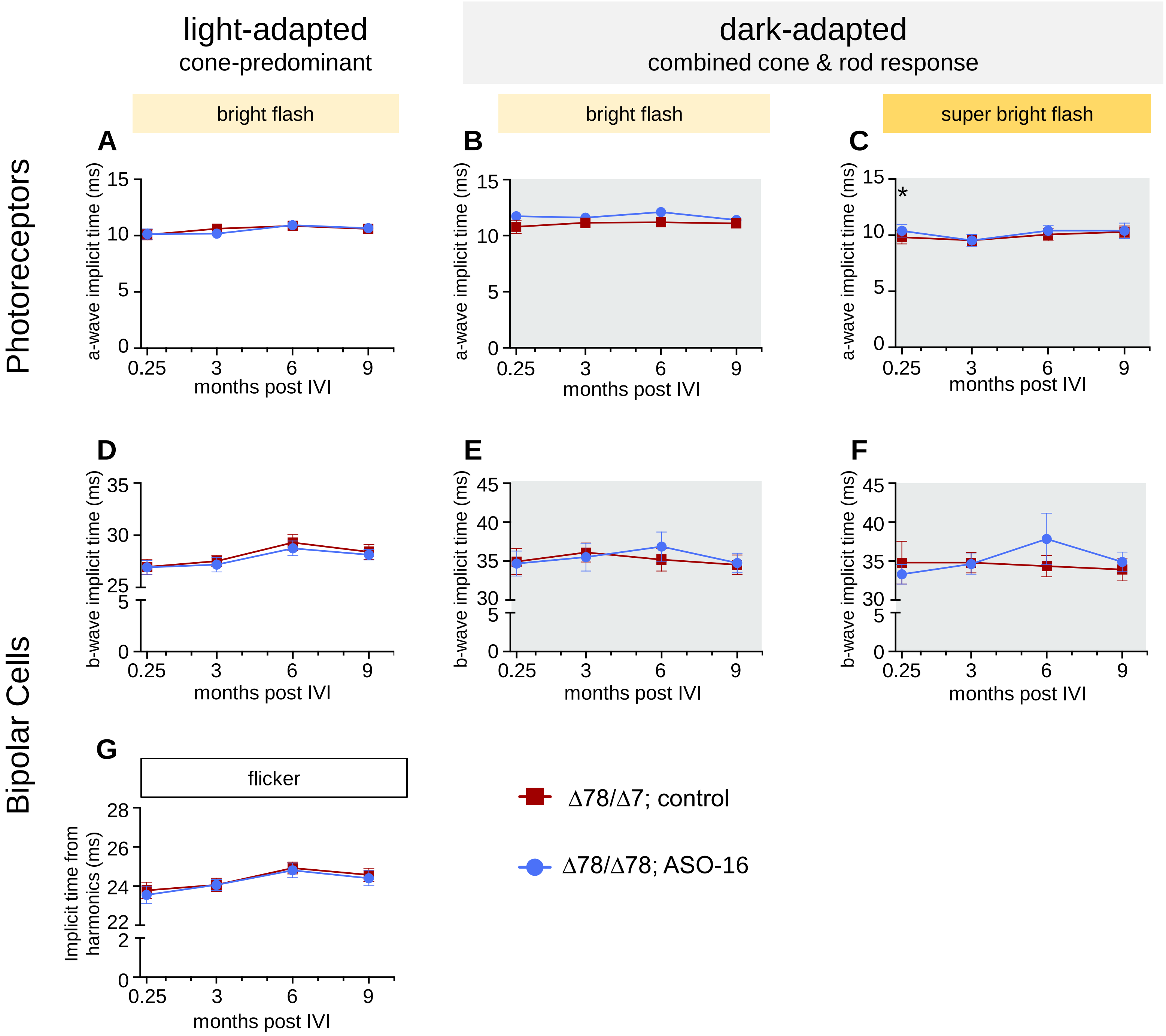
light-adapted
cone-predominant
dark-adapted
combined cone & rod response
bright flash
bright flash
super bright flash
Photoreceptors
Bipolar Cells
A
B
C
D
E
F
G
a-wave implicit time (ms)
b-wave implicit time (ms)
months post IVI
flicker
Implicit time from harmonics (ms)
Δ78/Δ7; control
Δ78/Δ78; ASO-16

**Figure S5**

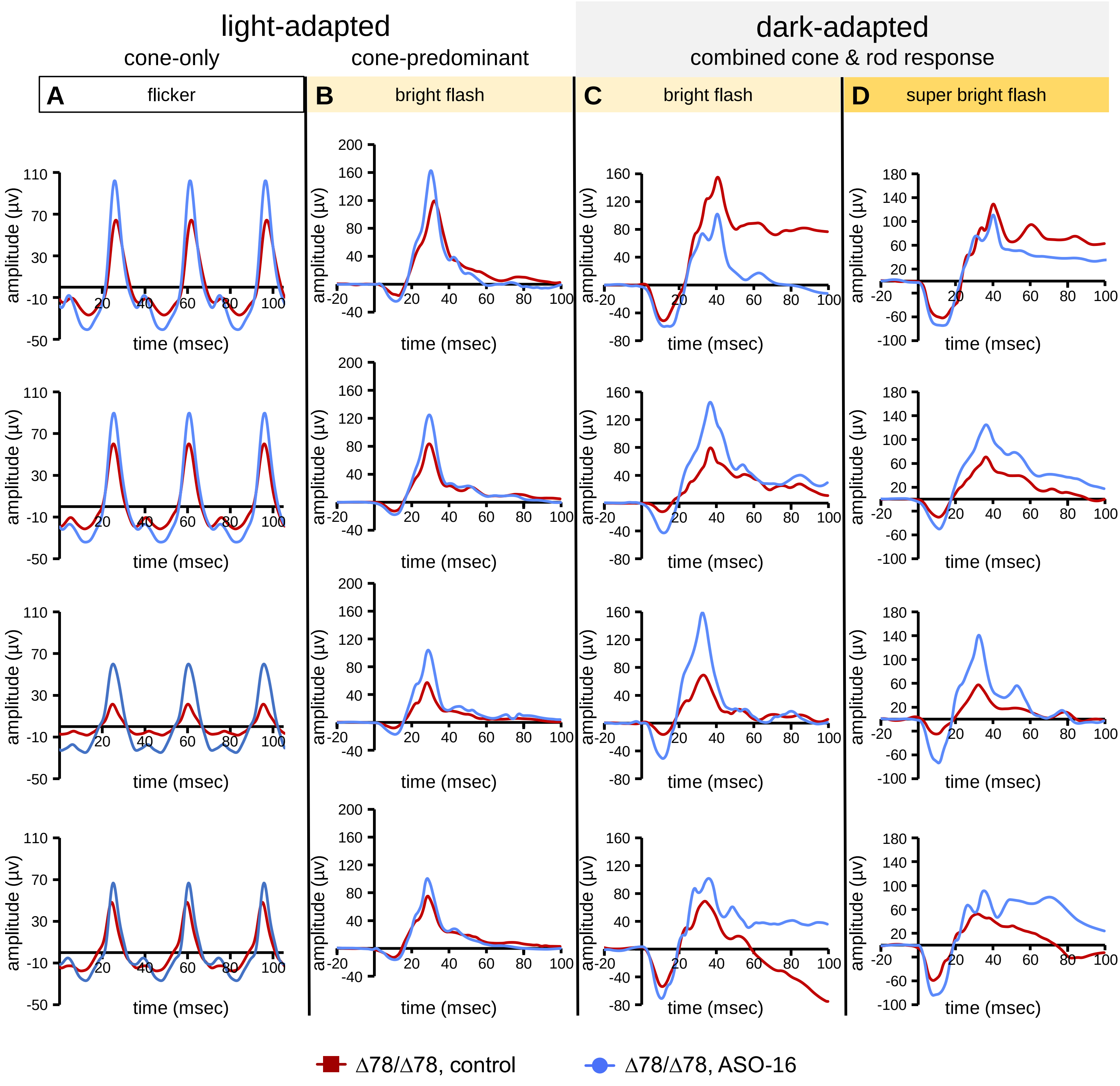
light-adapted
cone-only
cone-predominant
dark-adapted
combined cone & rod response
A
flicker
B
bright flash
C
bright flash
D
super bright flash
amplitude (µv)
time (msec)
Δ78/Δ78, control
Δ78/Δ78, ASO-16

**Figure S6**

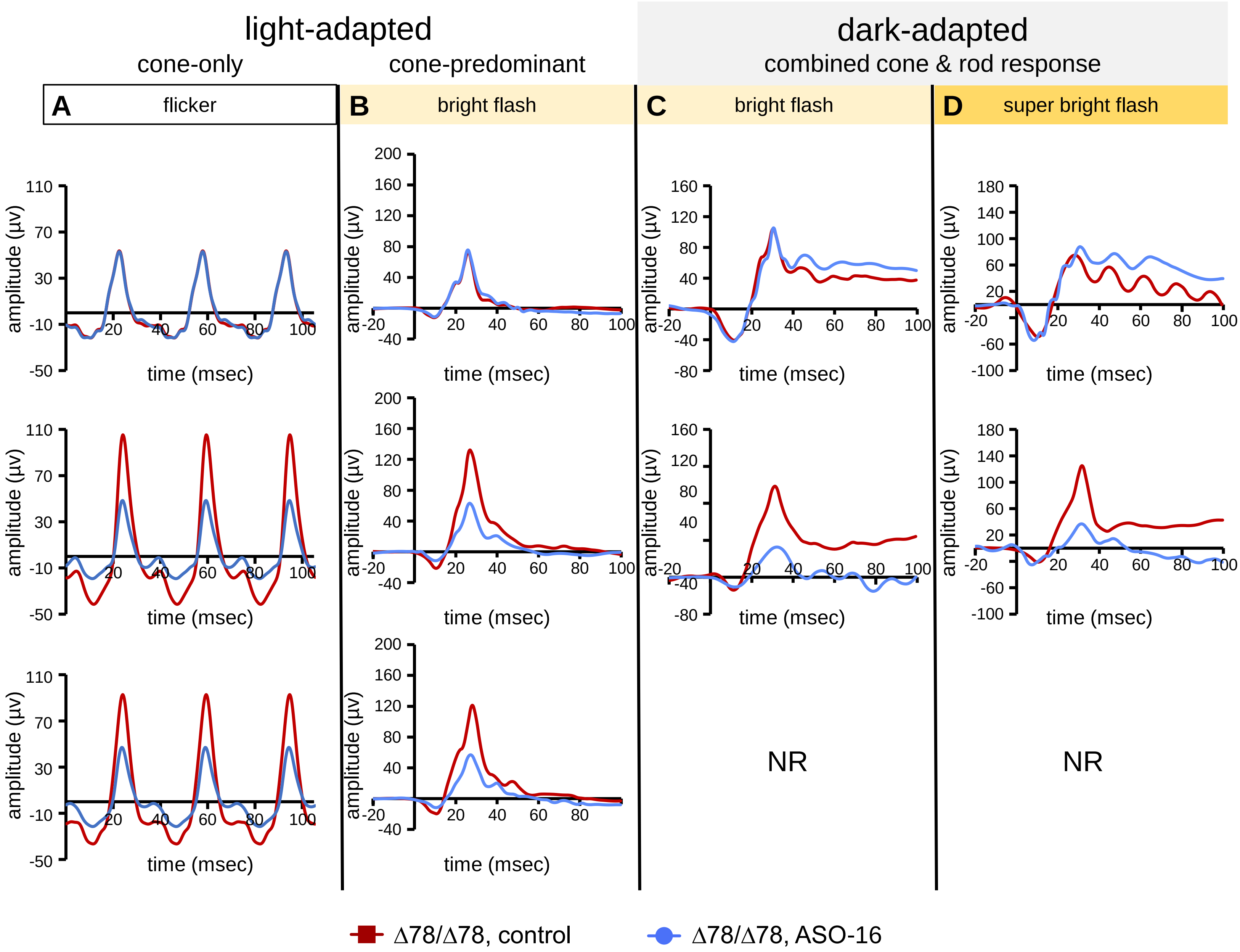
light-adapted
cone-only
cone-predominant
dark-adapted
combined cone & rod response
A
flicker
B
bright flash
C
bright flash
D
super bright flash
amplitude (μv)
time (msec)
NR
NR
Δ78/Δ78, control
Δ78/Δ78, ASO-16

**Figure S7**

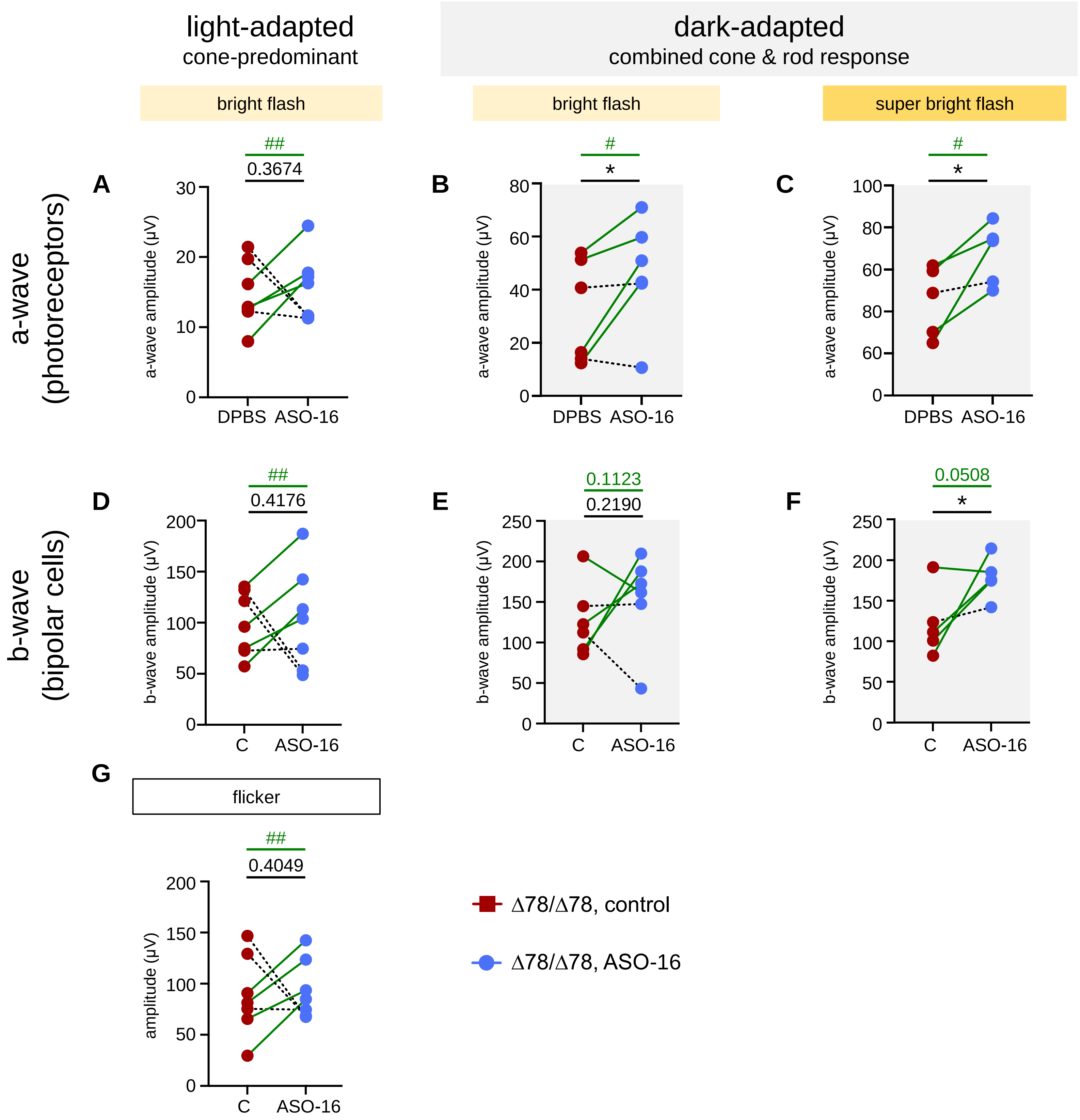

light-adapted
cone-predominant
dark-adapted
combined cone & rod response
bright flash
bright flash
super bright flash
a-wave
(photoreceptors)
b-wave
(bipolar cells)
A
B
C
D
E
F
G
##
0.3674
#
*
#
*
##
0.4176
0.1123
0.2190
0.0508
*
flicker
##
0.4049
a-wave amplitude (µV)
b-wave amplitude (µV)
amplitude (µV)
DPBS
ASO-16
C
ASO-16
Δ78/Δ78, control
Δ78/Δ78, ASO-16


**Figure S8**

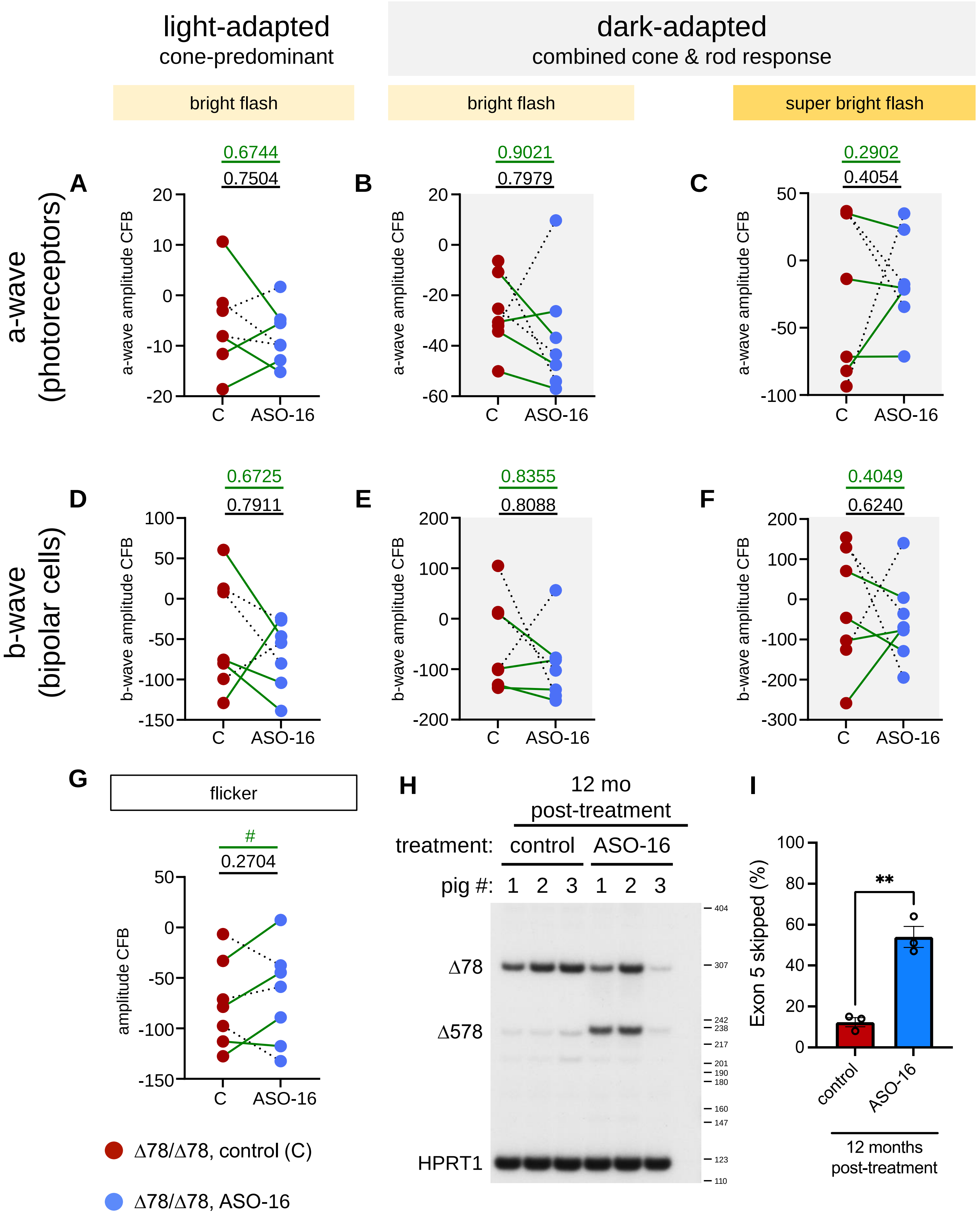

light-adapted
cone-predominant
dark-adapted
combined cone & rod response
bright flash
bright flash
super bright flash
a-wave
(photoreceptors)
b-wave
(bipolar cells)
A
0.6744
0.7504
a-wave amplitude CFB
C
ASO-16
B
0.9021
0.7979
C
0.2902
0.4054
D
0.6725
0.7911
b-wave amplitude CFB
E
0.8355
0.8088
F
0.4049
0.6240
G
flicker
#
0.2704
amplitude CFB
Δ78/Δ78, control (C)
Δ78/Δ78, ASO-16
H
12 mo
post-treatment
treatment: control ASO-16
pig #: 1 2 3 1 2 3
Δ78
Δ578
HPRT1
I
Exon 5 skipped (%)
**
control
ASO-16
12 months
post-treatment


**Figure S9**

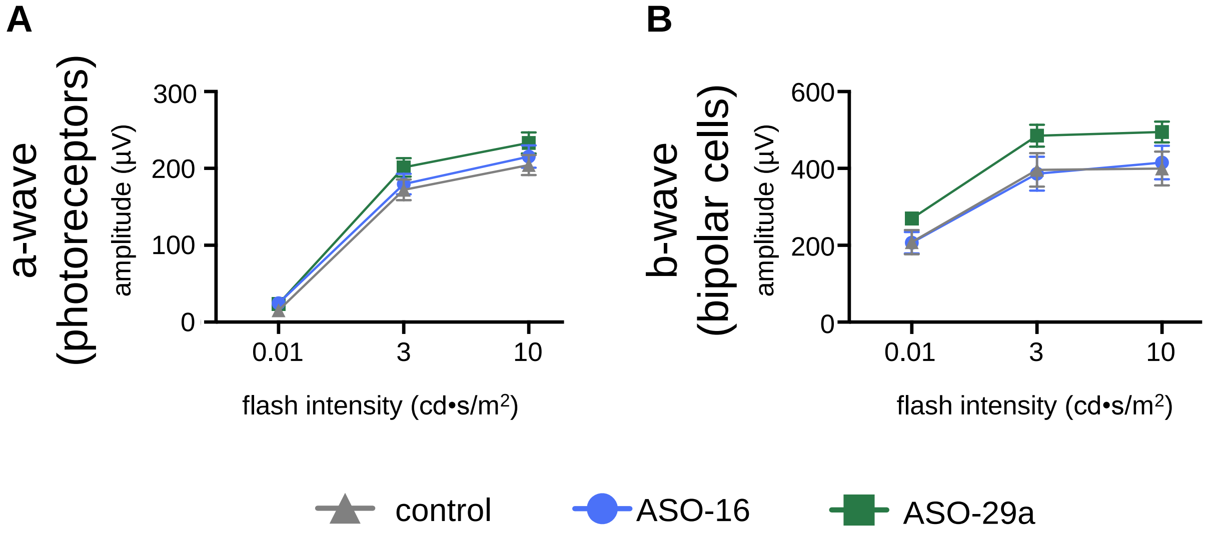
A
B
a-wave
(photoreceptors)
amplitude (μV)
b-wave
(bipolar cells)
300
200
100
0
600
400
200
0
0.01
3
10
flash intensity (cd•s/m2)
control
ASO-16
ASO-29a

**Figure S10**

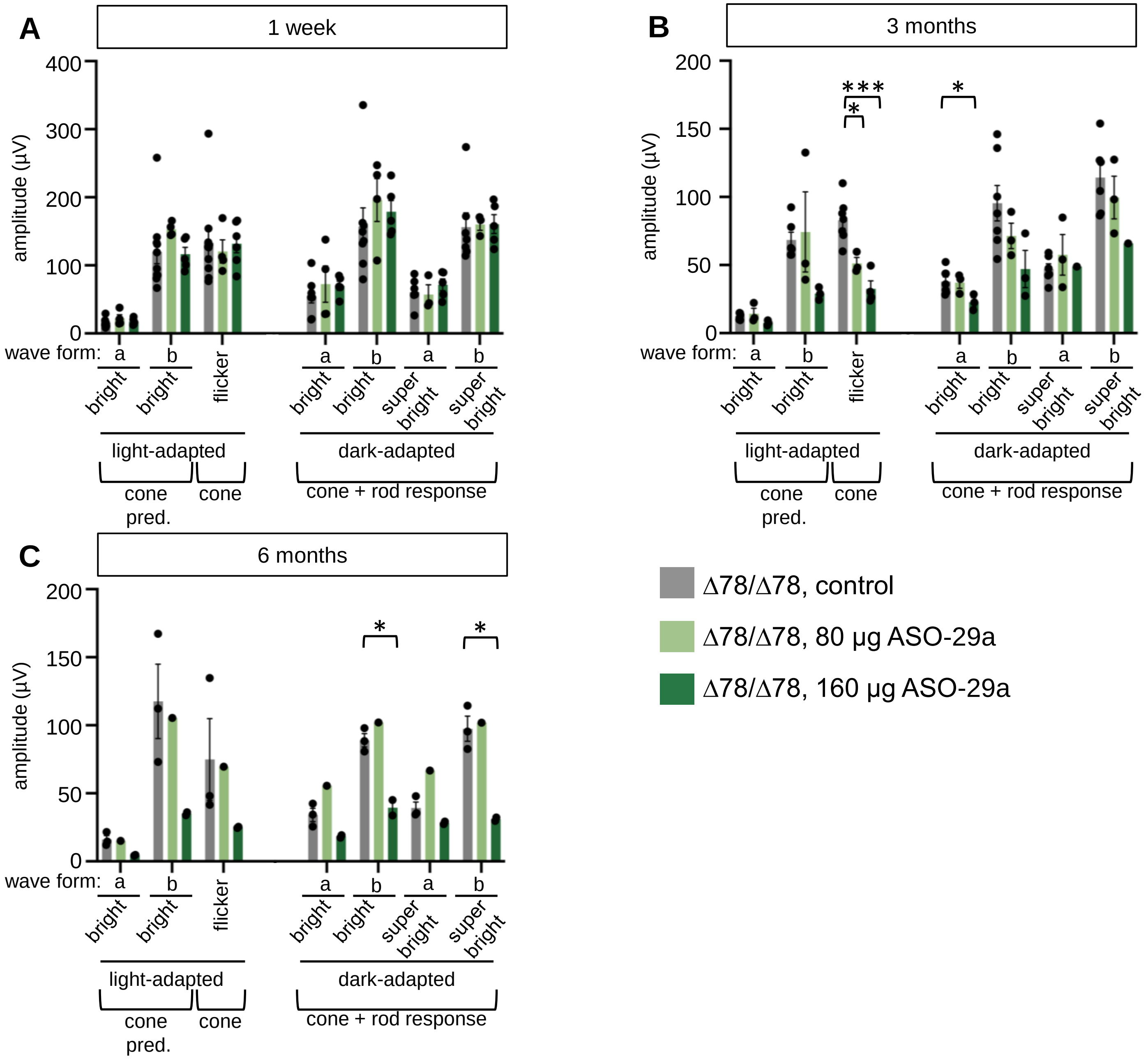
A
1 week
amplitude (µV)
400
300
200
100
0
wave form:
a
b
bright
bright
flicker
a
b
a
b
bright
bright
super bright
super bright
light-adapted
cone pred.
cone
dark-adapted
cone + rod response
B
3 months
amplitude (µV)
200
150
100
50
0
***
*
*
C
6 months
amplitude (µV)
200
150
100
50
0
*
*
Δ78/Δ78, control
Δ78/Δ78, 80 µg ASO-29a
Δ78/Δ78, 160 µg ASO-29a

**Figure S11**

**Table S1**. Ocular and Non-ocular Adverse Events

Summary of ocular AEs

**Figure 2**

| | 0.9% saline (n = 6) | DPBS (n = 3) | ASO-16, 300 µg (n = 2) | ASO-16, 200 µg (n = 4) | ASO-16, 80 µg (n = 3) | Relationship to study drug |
|---|---|---|---|---|---|---|
| Age at treatment (days) | 11-30 | 125-154 | 11-30 | 11-30 | 125-154 | |
| Number of eyes (%) | | | | | | |
| **Ocular AE** | **1 (16.7)** | **1 (33.3)** | **2 (100)** | **3 (75)** | **0 (0)** | |
| cataract | 1 (16.7) | 0 | 0 | 0 | 0 | not related |
| cloudy vitreous | 1 (16.7) | 0 | 0 | 0 | 0 | not related |
| corneal ulcer | 0 | 1 (33.3) | 0 | 0 | 0 | not related |
| diminished ERG | 0 | 0 | 2 (100) | 3 (75) | 0 | probable |
| **Non-ocular AE** | **0 (0)** | **0 (0)** | **0 (0)** | **4 (100)** | **0 (0)** | |
| (possible) influenza | 0 | 0 | 0 | 3 (75) | 0 | unlikely |
| GI issues resulting in humane endpoint due to weight loss | 0 | 0 | 0 | 1 (25) | 0 | unlikely |

**Figure 3; S4; S5**

| | DPBS (n = 15) | ASO-16, 80 µg (n = 9) | Relationship to study drug |
|---|---|---|---|
| Age at treatment (days) | 103-154 | 111-154 | |
| Number of eyes (%) | | | |
| **Ocular AE** | **1 (6.7)** | **0 (0)** | |
| cataract | 1 (6.7) | 0 | not related |
| **Non-ocular AE** | **2 (13.3)** | **1 (11.1)** | |
| (possible) meningitis | 1 (6.7) | 0 | unlikely |
| leg injury | 1 (6.7) | 1 (11.1) | unlikely |

**Figure 6; S11**

| | DPBS (n = 5) | ASO-25, 80 µg (n = 5) | ASO-25, 160 µg (n = 5) | Relationship to study drug |
|---|---|---|---|---|
| Age at treatment (days) | 103-178 | 103-178 | 103-118 | |
| Number of eyes (%) | | | | |
| **Ocular AE** | **1 (20)** | **0 (0)** | **1 (20)** | |
| cataract | 1 (20) | 0 | 1 (20) | possible |

**Table S2**. Figure 3 ERG p-values

| **Fig 3D: light-adapted, 8.0 cd•s/m$^2$ bright flash** | | p-value (sample number, N) | | |
|---|---|---|---|---|
| a-wave CFB | | 3 mo | 6 mo | 9 mo |
| paired t-test | Δ78/Δ78, C vs. Δ78/Δ78, ASO-16 (80 µg) | 0.0570 (7) | 0.3416 (7) | 0.2877 (7) |
| one-way ANOVA | +/Δ78, C vs. Δ78/Δ78, C | **0.0090** (6, 7) | 0.1566 (6, 7) | 0.0748 (6, 7) |
| | +/Δ78, C vs. Δ78/Δ78, ASO-16 (80 µg ) | 0.1357 (6, 7) | 0.2549 (6, 7) | 0.1982 (6, 7) |

| **Fig 3E: dark-adapted, 8.0 cd•s/m$^2$ bright flash** | | p-value | | |
|---|---|---|---|---|
| a-wave CFB | | 3 mo | 6 mo | 9 mo |
| paired t-test | Δ78/Δ78, C vs. Δ78/Δ78, ASO-16 (80 µg) | 0.6664 (6) | 0.9592 (6) | 0.0965 (6) |
| one-way ANOVA | +/Δ78, C vs. Δ78/Δ78, C | 0.4134 (6, 6) | 0.1665 (6, 7) | 0.2150 (6, 7) |
| | +/Δ78, C vs. Δ78/Δ78, ASO-16 (80 µg ) | 0.1858 (6, 6) | 0.0028 (6, 6) | 0.7254 (6, 6) |

| **Fig 3F: dark-adapted, 25.0 cd•s/m$^2$ bright flash** | | p-value | | |
|---|---|---|---|---|
| a-wave CFB | | 3 mo | 6 mo | 9 mo |
| paired t-test | Δ78/Δ78, C vs. Δ78/Δ78, ASO-16 (80 µg) | 0.2376 (7) | 0.3376 (5) | **0.0025** (5) |
| one-way ANOVA | +/Δ78, C vs. Δ78/Δ78, C | 0.8493 (6, 7) | 0.6126 (6, 7) | 0.1809 (6, 6) |
| | +/Δ78, C vs. Δ78/Δ78, ASO-16 (80 µg ) | 0.9961 (6, 7) | 0.9604 (6, 5) | 0.7098 (6, 5) |

| **Fig 3G: light-adapted, 8.0 cd•s/m$^2$ bright flash** | | p-value | | |
|---|---|---|---|---|
| b-wave CFB | | 3 mo | 6 mo | 9 mo |
| paired t-test | Δ78/Δ78, C vs. Δ78/Δ78, ASO-16 (80 µg) | **0.0062** (7) | **0.0074** (7) | **0.0497** (7) |
| one-way ANOVA | +/Δ78, C vs. Δ78/Δ78, C | **0.0251** (6, 7) | 0.7795 (6, 7) | **0.0359** (6, 7) |
| | +/Δ78, C vs. Δ78/Δ78, ASO-16 (80 µg ) | 0.6512 (6, 7) | 0.6435 (6, 7) | 0.8037 (6, 7) |

| **Fig 3H: dark-adapted, 8.0 cd•s/m$^2$ bright flash** | | p-value | | |
|---|---|---|---|---|
| b-wave CFB | | 3 mo | 6 mo | 9 mo |
| paired t-test | Δ78/Δ78, C vs. Δ78/Δ78, ASO-16 (80 µg) | 0.3827 (6) | 0.7515 (6) | 0.2182 (6) |
| one-way ANOVA | +/Δ78, C vs. Δ78/Δ78, C | 0.1107 (6, 6) | **0.0023** (6, 7) | **0.0367** (6, 7) |
| | +/Δ78, C vs. Δ78/Δ78, ASO-16 (80 µg ) | 0.2114 (6, 6) | **0.0006** (6, 6) | 0.2022 (6, 6) |

| **Fig 3I: dark-adapted, 25.0 cd•s/m$^2$ bright flash** | | p-value | | |
|---|---|---|---|---|
| b-wave CFB | | 3 mo | 6 mo | 9 mo |
| paired t-test | Δ78/Δ78, C vs. Δ78/Δ78, ASO-16 (80 µg) | 0.1892 (7) | 0.2696 (5) | **<0.0001** (5) |
| one-way ANOVA | +/Δ78, C vs. Δ78/Δ78, C | 0.6045 (6, 7) | 0.1700 (6, 7) | 0.1488 (6, 6) |
| | +/Δ78, C vs. Δ78/Δ78, ASO-16 (80 µg ) | 0.9940 (6, 7) | 0.4226 (6, 5) | 0.8304 (6, 5) |

| **Fig 3K: 8.0 cd•s/m$^2$ flicker at 28.3 Hz** | | p-value | | |
|---|---|---|---|---|
| amplitude CFB | | 3 mo | 6 mo | 9 mo |
| paired t-test | Δ78/Δ78, C vs. Δ78/Δ78, ASO-16 (80 µg) | **0.0183** (7) | 0.1218 (7) | 0.0898 (7) |
| one-way ANOVA | +/Δ78, C vs. Δ78/Δ78, C | **0.0032** (6, 7) | 0.1443 (6, 7) | **0.0218** (5, 7) |
| | +/Δ78, C vs. Δ78/Δ78, ASO-16 (80 µg ) | 0.6268 (6, 7) | 0.7008 (6, 7) | 0.4200 (5, 7) |

Sample size (N) is indicated in parenthesis next to p-value for each comparison. In some cases, a measurement is missing due to technical issues as described in methods.

**Table S3**. ERG p-values corresponding to results from Figure 4

| **Fig 4A: light-adapted, 8.0 cd•s/m$^2$ bright flash** | | p-value (sample number, N) |
|---|---|---|
| a-wave CFB | | 9 mo |
| paired t-test | Δ78/Δ78: C vs. ASO-16 (80 µg), high responders | 0.0958 (4) |
| paired t-test | Δ78/Δ78: C vs. ASO-16 (80 µg), all animals | 0.2877 (7) |

| **Fig 4B: dark-adapted, 8.0 cd•s/m$^2$ bright flash** | | p-value |
|---|---|---|
| a-wave CFB | | 9 mo |
| paired t-test | Δ78/Δ78: C vs. ASO-16 (80 µg), high responders | 0.0322 (4) |
| paired t-test | Δ78/Δ78: C vs. ASO-16 (80 µg), all animals | 0.0965 (6) |

| **Fig 4C: dark-adapted, 25.0 cd•s/m$^2$ bright flash** | | p-value |
|---|---|---|
| a-wave CFB | | 9 mo |
| paired t-test | Δ78/Δ78: C vs. ASO-16 (80 µg), high responders | **0.0049** (4) |
| paired t-test | Δ78/Δ78: C vs. ASO-16 (80 µg), all animals | **0.0025** (5) |

| **Fig 4D: light-adapted, 8.0 cd•s/m$^2$ bright flash** | | p-value |
|---|---|---|
| b-wave CFB | | 9 mo |
| paired t-test | Δ78/Δ78: C vs. ASO-16 (80 µg), high responders | **0.0068** (4) |
| paired t-test | Δ78/Δ78: C vs. ASO-16 (80 µg), all animals | **0.0497** (7) |

| **Fig 4E: dark-adapted, 8.0 cd•s/m$^2$ bright flash** | | p-value |
|---|---|---|
| b-wave CFB | | 9 mo |
| paired t-test | Δ78/Δ78: C vs. ASO-16 (80 µg), high responders | 0.1120 (4) |
| paired t-test | Δ78/Δ78: C vs. ASO-16 (80 µg), all animals | 0.2182 (6) |

| **Fig 4F: dark-adapted, 25.0 cd•s/m$^2$ bright flash** | | p-value |
|---|---|---|
| b-wave CFB | | 9 mo |
| paired t-test | Δ78/Δ78: C vs. ASO-16 (80 µg), high responders | **0.0009** (4) |
| paired t-test | Δ78/Δ78: C vs. ASO-16 (80 µg), all animals | **<0.0001** (5) |

| **Fig 4G: 8.0 cd•s/m$^2$ flicker at 28.3 Hz** | | p-value |
|---|---|---|
| amplitude CFB | | 9 mo |
| paired t-test | Δ78/Δ78: C vs. ASO-16 (80 µg), high responders | **0.0106** (4) |
| paired t-test | Δ78/Δ78: C vs. ASO-16 (80 µg), all animals | 0.0898 (7) |

Sample size (N) is indicated in parenthesis next to p-value for each comparison. In some cases, a measurement is missing due to technical issues as described in methods.

**Table S4**. ERG p-values corresponding to results from Figure S8

| **Fig S8A: light-adapted, 8.0 cd•s/m$^2$ bright flash** | | p-value (sample number, N) |
|---|---|---|
| a-wave CFB | | 9 mo |
| paired t-test | Δ78/Δ78: C vs. ASO-16 (80 μg), high responders | **0.0086** (4) |
| paired t-test | Δ78/Δ78: C vs. ASO-16 (80 μg), all animals | 0.3674 (7) |

| **Fig S8B: dark-adapted, 8.0 cd•s/m$^2$ bright flash** | | p-value |
|---|---|---|
| a-wave CFB | | 9 mo |
| paired t-test | Δ78/Δ78: C vs. ASO-16 (80 μg), high responders | **0.0166** (4) |
| paired t-test | Δ78/Δ78: C vs. ASO-16 (80 μg), all animals | **0.0325** (6) |

| **Fig S8C: dark-adapted, 25.0 cd•s/m$^2$ bright flash** | | p-value |
|---|---|---|
| a-wave CFB | | 9 mo |
| paired t-test | Δ78/Δ78: C vs. ASO-16 (80 μg), high responders | **0.0209** (4) |
| paired t-test | Δ78/Δ78: C vs. ASO-16 (80 μg), all animals | **0.0192** (5) |

| **Fig S8D: light-adapted, 8.0 cd•s/m$^2$ bright flash** | | p-value |
|---|---|---|
| b-wave CFB | | 9 mo |
| paired t-test | Δ78/Δ78: C vs. ASO-16 (80 μg), high responders | **0.0023** (4) |
| paired t-test | Δ78/Δ78: C vs. ASO-16 (80 μg), all animals | 0.4176 (7) |

| **Fig S8E: dark-adapted, 8.0 cd•s/m$^2$ bright flash** | | p-value |
|---|---|---|
| b-wave CFB | | 9 mo |
| paired t-test | Δ78/Δ78: C vs. ASO-16 (80 μg), high responders | 0.1123 (4) |
| paired t-test | Δ78/Δ78: C vs. ASO-16 (80 μg), all animals | 0.2190 (6) |

| **Fig S8F: dark-adapted, 25.0 cd•s/m$^2$ bright flash** | | p-value |
|---|---|---|
| b-wave CFB | | 9 mo |
| paired t-test | Δ78/Δ78: C vs. ASO-16 (80 μg), high responders | 0.0508 (4) |
| paired t-test | Δ78/Δ78: C vs. ASO-16 (80 μg), all animals | **0.0386** (5) |

| **Fig S8G: 8.0 cd•s/m$^2$ flicker at 28.3 Hz** | | p-value |
|---|---|---|
| amplitude CFB | | 9 mo |
| paired t-test | Δ78/Δ78: C vs. ASO-16 (80 μg), high responders | **0.0027** (4) |
| paired t-test | Δ78/Δ78: C vs. ASO-16 (80 μg), all animals | 0.4049 (7) |

Sample size (N) is indicated in parenthesis next to p-value for each comparison. In some cases, a measurement is missing due to technical issues as described in methods.

**Table S5**. ERG p-values corresponding to results from Figure S9

| **Fig S9A: light-adapted, 8.0 cd•s/m$^2$ bright flash** | | p-value |
|---|---|---|
| a-wave CFB | | 12 mo |
| paired t-test | DPBS Δ78/Δ78 vs. ASO-16, 80 µg Δ78/Δ78, high responders | 0.6744 (4) |
| paired t-test | DPBS Δ78/Δ78 vs. ASO-16, 80 µg Δ78/Δ78, all animals | 0.7504 (7) |

| **Fig S9B: dark-adapted, 8.0 cd•s/m$^2$ bright flash** | | p-value |
|---|---|---|
| a-wave CFB | | 12 mo |
| paired t-test | DPBS Δ78/Δ78 vs. ASO-16, 80 µg Δ78/Δ78, high responders | 0.9021 (4) |
| paired t-test | DPBS Δ78/Δ78 vs. ASO-16, 80 µg Δ78/Δ78, all animals | 0.7979 (7) |

| **Fig S9C: dark-adapted, 25.0 cd•s/m$^2$ bright flash** | | p-value |
|---|---|---|
| a-wave CFB | | 12 mo |
| paired t-test | DPBS Δ78/Δ78 vs. ASO-16, 80 µg Δ78/Δ78, high responders | 0.2902 (4) |
| paired t-test | DPBS Δ78/Δ78 vs. ASO-16, 80 µg Δ78/Δ78, all animals | 0.4054 (7) |

| **Fig S9D: light-adapted, 8.0 cd•s/m$^2$ bright flash** | | p-value |
|---|---|---|
| b-wave CFB | | 12 mo |
| paired t-test | DPBS Δ78/Δ78 vs. ASO-16, 80 µg Δ78/Δ78, high responders | 0.6725 (4) |
| paired t-test | DPBS Δ78/Δ78 vs. ASO-16, 80 µg Δ78/Δ78, all animals | 0.7911 (7) |

| **Fig S9E: dark-adapted, 8.0 cd•s/m$^2$ bright flash** | | p-value |
|---|---|---|
| b-wave CFB | | 12 mo |
| paired t-test | DPBS Δ78/Δ78 vs. ASO-16, 80 µg Δ78/Δ78, high responders | 0.8355 (4) |
| paired t-test | DPBS Δ78/Δ78 vs. ASO-16, 80 µg Δ78/Δ78, all animals | 0.8088 (7) |

| **Fig S9F: dark-adapted, 25.0 cd•s/m$^2$ bright flash** | | p-value |
|---|---|---|
| b-wave CFB | | 12 mo |
| paired t-test | DPBS Δ78/Δ78 vs. ASO-16, 80 µg Δ78/Δ78, high responders | 0.4049 (4) |
| paired t-test | DPBS Δ78/Δ78 vs. ASO-16, 80 µg Δ78/Δ78, all animals | 0.6240 (7) |

| **Fig S9G: 8.0 cd•s/m$^2$ flicker at 28.3 Hz** | | p-value |
|---|---|---|
| amplitude | | 12 mo |
| paired t-test | DPBS Δ78/Δ78 vs. ASO-16, 80 µg Δ78/Δ78, high responders | **0.0423** (4) |
| paired t-test | DPBS Δ78/Δ78 vs. ASO-16, 80 µg Δ78/Δ78, all animals | 0.2704 (7) |

**Table S6**. Splice switching antisense oligonucleotide and primer sequences

| ASOs | Sequence (5'-3') |
|---|---|
| 1 | CAGAGAACACAGTGAGAC |
| 2 | GGGACCAGAGAACACAGT |
| 3 | CGCCTGGGACCAGAGAAC |
| 4 | AGCACCGCCTGGGACCAG |
| 5 | CCAGGAGCACCGCCTGGG |
| 6 | GTCTGCCAGGAGCACCGC |
| 7 | AGGATGTCTGCCAGGAGC |
| 8 | TGGGAAGGATGTCTGCCA |
| 9 | GAGGGTGGGAAGGATGTC |
| 10 | ATGATGAGGGTGGGAAGG |
| 11 | ATTTGATGATGAGGGTGG |
| 12 | CAGCAATTTGATGATGAG |
| 13 | GGAGCCAGCAATTTGATG |
| 14 | CAAGAGGAGCCAGCAATT |
| 15 | GAGGCCAAGAGGAGCCAG |
| 16 | AGATGGAGGCCAAGAGGA |
| 17 | GCAGCAGATGGAGGCCAA |
| 18 | GTAGGGCAGCAGATGGAG |
| 19 | GACCTGTAGGGCAGCAGA |
| 20 | CCCCAGACCTGTAGGGCA |
| 21 | CCCCTCCCCAGACCTGTA |
| 22 | CCCACCCCCTCCCCAGAC |
| 23 | CCACCCCCACCCCCTCCC |
| 24 | TCCCACCCCCACCCCCTC |
| 29a | CCCAGACCTGTAGGGCAG |
| 29b | CCAGACCTGTAGGGCAGC |
| 29c | CAGACCTGTAGGGCAGCA |
| 29d | AGACCTGTAGGGCAGCAG |
| **Primers** | **Sequence (5'-3')** |
| pCLN3ex3F | AGTGCTGCCCACGACATC |
| pCLN3ex4F | TAACTCTGTCTCCACGGC |
| pCLN3 ex6R | TTCCAGCGGCACAGATCC |
| pCLN3in6F | TGGTTTGCTAACTGGGTGGG |
| pCLN3in8R | TCACCACATGGAGAAGGAAGC |
| pCLN3ex10R | ACAGGTGTGAGCCGGAGC |
| pHPRT1F | TTATGGACAGGACTGAACGGC |
| pHPRT1R | GTAATCCAGCAGGTCAGCAAAG |
| hCLN3 ex4 | GCAACTCTGTCTCTACGGC |
| hCLN3 ex10 | CTTGAACACTGTCCACC |

**A**

**Genomic matches of ASO-29a subsequences**

| Length (nt) | Number of possible subsequences | Number of off-target hits | Off-target locus (hg19) | Gene annotation (RefSeq) | | |
|---|---|---|---|---|---|---|
| | | | | Gene | Strand | Region |
| 18 | 1 | 0 | - | - | - | - |
| 17 | 2 | 0 | - | - | - | - |
| 16 | 3 | 6 | chr1:970996-971013 | *AGRN* | sense | intron (291 bp away from junction) |
| | | | chr6:1859273-1859290 | *GMDS* | antisense | intron (>71kb away from junction) |
| | | | chr11:117908797-117908814 | *SMIM35* | sense | intron (>6kb away from junction) |
| | | | chr13:55752072-55752089 | - | - | - |
| | | | chr15:86917573-86917590 | *AGBL1* | antisense | intron (>23kb away from junction) |
| | | | chr17:72822712-72822729 | *TMEM104* | antisense | intron (>6.7kb away from junction) |

**B**

**Genomic matches of nusinersen subsequences**

| Length (nt) | Number of possible subsequences | Number of off-target hits | Off-target locus (hg19) | Gene annotation (RefSeq) | | |
|---|---|---|---|---|---|---|
| | | | | Gene | Strand | Region |
| 18 | 1 | 0 | - | - | - | - |
| 17 | 2 | 3 | chr2: 151151798-151151814 | - | - | - |
| | | | chr6: 137801026-137801042 | - | - | - |
| | | | chr12: 129939692-129939708 | *TMEM132D* | sense | intron (>75 kb away from junction) |
| 16 | 3 | 7 | chr13: 30033807-30033822 | *MTUS2* | antisense | intron (>19kb away from junction) |
| | | | chr3: 82606750-82606765 | - | - | - |
| | | | chr15: 32439151-32439166 | *CHRNA7* | antisense | intron (>6.9kb away from junction) |
| | | | chr20: 9638297-9638312 | *PAK5* | sense | intron (>13kb away from junction) |
| | | | chr7: 125728305-125728320 | - | - | - |
| | | | chr7: 141917297-141917312 | *MGAM2* | antisense | intron (>800bp away from junction) |
| | | | chr3: 110103460-110103475 | - | - | - |

**Table S7. Genomic off-target footprints of ASO-29a, compared to those of nusinersen.** (**A**) All possible coding subsequences of a given length were taken from the ASO-29a sequence and aligned to the reference human genome in the *GRCh37/hg19* database. The number and identity of the off-target matches are shown. If the off-target locus is in an intron, the distance from the closest splice junction is indicated. (**B**) The same alignment was performed as in panel A, but for nusinersen in the human genome.

**Table S8**. Experimental Sample Sizes

| **Figure 2** | | | | | | | | | | | | | | | | |
|---|---|---|---|---|---|---|---|---|---|---|---|---|---|---|---|---|
| ASO Dose | vehicle | | 300 µg | | vehicle | | | 200 µg | | | vehicle | | | 80 µg | | |
| Time post-IVI | 3 mo | 6 mo | 3 mo | 6 mo | 3 mo | 6 mo | 12 mo | 3 mo | 6 mo | 12 mo | 3 mo | 6 mo | 12 mo | 3 mo | 6 mo | 12 mo |
| Sample Size (# of eyes) | 1 | 1 | 1 | 1 | 1 | 1 | 1 | 1 | 1 | 1 | 1 | 1 | 1 | 1 | 1 | 1 |

| **Figure 3, S4, S5** | | | |
|---|---|---|---|
| Time post-IVI | DPBS +/Δ78 | DPBS Δ78/Δ78 | ASO-16, 80 µg Δ78/Δ78 |
| 1 week | 7 (5M, 2F) | 9 (6M, 3F) | 9 (6M, 3F) |
| 3 months | 7 (5M, 2F) | 9 (6M, 3F) | 9 (6M, 3F) |
| 6 months | 7 (5M, 2F) | 9 (6M, 3F) | 9 (6M, 3F) |
| 9 months | 7 (5M, 2F) | 9 (6M, 3F) | 9 (6M, 3F) |

| **Figure 6** | | | | | | | | | | | | |
|---|---|---|---|---|---|---|---|---|---|---|---|---|
| ASO Dose | vehicle | | | 80 µg | | | vehicle | | | 160 µg | | |
| Time post-IVI | 1 mo | 3 mo | 6 mo | 1 mo | 3 mo | 6 mo | 1 mo | 3 mo | 6 mo | 1 mo | 3 mo | 6 mo |
| Sample Size (# of eyes) | 2 | 2 | 1 | 2 | 2 | 1 | 1 | 2 | 2 | 1 | 2 | 2 |

| **Figure S3** | | | | | | | | |
|---|---|---|---|---|---|---|---|---|
| Genotype | WT | | | | Δ78 | | | |
| Age | 2 | 6 | 36 | 50 | 2 | 6 | 36 | 50 |
| Sample Size | 2 | 1 | 1 | 1 | 2 | 2 | 3 | 2 |

| **Figure S10** | | | |
|---|---|---|---|
| Treatment | DPBS | ASO-16 | ASO-29a |
| Genotype | WT | WT | WT |
| Sample Size (# of eyes) | 4 | 8 | 8 |

| **Figure S11** | | | |
|---|---|---|---|
| Time post-IVI | DPBS Δ78/Δ78 | ASO-29a, 80 µg Δ78/Δ78 | ASO-29a, 160 µg Δ78/Δ78 |
| 1 week | 10 (4M, 6F) | 4 (2M, 2F) | 5 (2M, 3F) |
| 3 months | 6 (4M, 2F) | 3 (3M, 0F) | 3 (2M, 1F) |
| 6 months | 3 (2M, 1F) | 1 (1M, 0F) | 2 (1M, 1F) |
